# Transcriptomic response to divergent selection for flowering times reveals convergence and key players of the underlying gene regulatory network

**DOI:** 10.1101/461947

**Authors:** Maud I. Tenaillon, Khawla Seddiki, Maeva Mollion, Martine Le Guilloux, Elodie Marchadier, Adrienne Ressayre, Christine Dillmann

**Affiliations:** UMR GQE-Le Moulon, INRA-UPS-CNRS-AgroParisTech, Université Paris-Saclay – Gif-sur-Yvette, France

**Keywords:** Floral Transition, Gene network, Flowering, Maize, Response to selection, Convergence, RNA-seq

## Abstract

Artificial selection experiments are designed to investigate phenotypic evolution of complex traits and its genetic basis. Here we focused on flowering time, a trait of key importance for plant adaptation and life-cycle shifts. We undertook divergent selection experiments from two maize inbred lines. After 13 generations of selection, we obtained a time-lag of roughly two weeks between Early- and Late-populations. We used this material to characterize the genome-wide transcriptomic response to selection in the shoot apical meristem before, during and after floral transition in field conditions during two consecutive years. We validated the reliability of performing RNA-sequencing in uncontrolled conditions. We found that roughly half of maize genes were expressed in the shoot apical meristem, 59.3% of which were differentially expressed. We detected a majority of genes with differential expression between inbreds and across meristem status, and retrieved a subset of 2,451 genes involved in the response to selection. Among these, we found a significant enrichment for genes with known function in maize flowering time. Furthermore, they were more often shared between inbreds than expected by chance, suggesting convergence of gene expression. We discuss new insights into the expression pattern of key players of the underlying gene regulatory network including the *Zea mays* genes *CENTRORADIALIS* (*ZCN8*), *RELATED TO AP2.7* (*RAP2.7*), *MADS4* (*ZMM4*), *KNOTTED1* (*KN1*), *GIBBERELLIN2*-*OXIDASE1* (*GA2ox1*), as well as alternative scenarios for genetic convergence.

## Introduction

Artificial selection experiments are designed to investigate evolutionary responses of complex traits. These responses inform us about the limits to the evolution of ecologically relevant phenotypes as well as their genetic architecture, determined by interactions among a multitude of environmentally sensitive genes. Replication of such experiments across distinct genetic backgrounds provides a unique opportunity to test whether convergent evolution at the phenotypic level recruits similar molecular solutions. Because artificial selection experiments are extremely time costly, they have most often been conducted in model organisms with a short generation time, prokaryotes such as *E. coli* (Tenaillon, et al. 2012; Good, et al. 2017) or, eukaryotes such as yeast (Burke, et al. 2014), fruitfly (Burke, et al. 2010; Graves, et al. 2017), domestic mouse (Chan, et al. 2012), *C. elegans* (McGrath, et al. 2011). These studies have collectively recovered genetic predictability at different levels, SNPs, genes and functional units. This predictability however depends on the origins of mutations that contribute to the response to selection. Hence, standing genetic variants shared between genetic backgrounds – inherited from a common ancestor – are more likely to generate convergence than unshared *de novo* mutations (Graves, et al. 2017).

Long-lasting evolution experiments are much rarer in vascular plants (but see Goldringer, et al. (2006), Roels and Kelly (2011), Gervasi and Schiestl (2017)). Maize however stands as an exception: Cyril G. Hopkins started the historical heritage of selection experiments in this model species by launching the Illinois divergent selection for protein and oil content in 1896 (Dudley and Lambert 1992). Since then, six other experiments have been undertaken: two for increased prolificacy (Maita and Coors 1996) and grain yield (Lamkey 1992), two for divergent seed size (Moose, et al. 2004) and ear length (Lopez-Reynoso and Hallauer 1998), and two for divergent flowering time (Durand, et al. 2010) named thereafter the Saclay divergent selection experiments. One important observation from these experiments is that the response to selection is in general, continuous over generations (reviewed in Lorant, et al. (2018)). This is particularly intriguing for the two Saclay divergent selection experiments that were conducted independently in two genetic backgrounds (two inbred lines) with limited standing variation (<1.9% residual heterozygosity) and very small population size. In these experiments, by applying divergent selection over 16 generations, we generated considerable phenotypic response with up to three-weeks difference between early and late flowering populations (Durand, et al. 2015), a range comparable to what is observed among the maize European Nested Association Mapping panel when evaluated across multiple environments (Lehermeier, et al. 2014). The dynamics of the response to selection in Saclay divergent selection experiments is consistent with a continuous input of new mutations (Durand, et al. 2010). The use of markers also revealed the contribution of complete sweeps from standing genetic variation (Durand, et al. 2015). The contribution of both, new mutations and standing variants, to the response to selection to flowering time is consistent with the known complexity of this trait – its high mutation target, i.e. >100 loci (Buckler, et al. 2009).

In plants, flowering is initiated by the transition of the shoot apical meristem from a vegetative status where the shoot apical meristem produces leaves, to a reproductive status where the shoot apical meristem produces reproductive organs. Because floral transition induces irreversible developmental changes that ultimately determine flowering time and the completion of seed development in suitable conditions, it is of key adaptive value. It is tuned by a gene regulatory network that integrates environmental and endogenous cues, and translates them to initiate flowering when the time is most favorable. This network is now well described in the model species *Arabidopsis thaliana* with few hundreds of described genes (Bouche, et al. 2016), but profound differences with maize have been pointed out. For instance in contrast to *A. thaliana*, maize does not exhibit vernalization response; and some of the major floral maize genes such as *Zea mays MADS4* (*ZMM4*) and *INDETERMINATE GROWTH1* (*ID1*) have no homologs in *A. thaliana* (Colasanti, et al. 1998).

In maize, the currently described gene regulatory network is still very limited. It encompasses 30 genes. Among them, *Zea mays CENTRORADIALIS* (*ZCN8*) encodes a florigen protein that migrates through the phloem from the leaf to the shoot apical meristem triggering via its accumulation, the reprogramming of the shoot apical meristem to floral transition (Meng, et al. 2011). *ZCN8* interacts with the floral activator *DELAYED FLOWERING1* (*DLF1*) (Muszynski, et al. 2006), and its expression is partially controlled by another activator, *ID1* (Meng, et al. 2011). *ZCN8* and *DLF1* act upstream *ZMM4*, a floral meristem identity integrator, which when overexpressed in the shoot apical meristem leads to early-flowering (Danilevskaya, et al. 2008). The transcription factor *RELATED TO AP2.7* (*RAP2.7*) encoded by a gene downstream of the cis-regulatory element *VGT1*, is a negative regulator of flowering time (Salvi, et al. 2007) and putatively modulates the expression of *ZMM4* (Dong, et al. 2012). Among other genes of the maize flowering pathway, *Zea mays CO-LIKE TIMING OF CAB1 PROTEIN DOMAIN* (*ZmCCT*) is central to photoperiod response (Hung, et al. 2012). Endogenous signals are delivered by the GA signaling pathway, the autonomous pathway, and the aging pathway through the action of miR156/miR172 genes. Interestingly, sucrose levels in sources leaves as well as carbohydrates export to sink tissues, also appear to play a main role in floral induction. For instance, metabolic signatures in mature leaves associate with the expression of *ID1*, and may contribute to the control of florigens (Coneva, et al. 2012).

Here we used early- and late-evolved genotypes from the two Saclay divergent selection experiments (Durand, et al. 2010), i.e. two different genetic backgrounds, to (i) characterize the genome-wide transcriptomic response to selection in the shoot apical meristem before, during and after floral transition; (ii) identify key players of the underlying gene regulatory; (iii) test for convergence of selection response at the gene level between the two backgrounds. We performed all our experiments on plants grown under agronomical field conditions.

## Results

We created a unique genetic material by applying divergent selection for flowering time independently to two maize inbred lines (F252 and MBS847-thereafter MBS) (Durand, et al. 2015). This material is used to investigate evolutionary responses to selection of a complex trait and to dissect its genetic architecture. After 13 generations of selection, phenotypic responses revealed roughly two weeks difference between Early and VeryLate F252 populations, and between Early and Late MBS populations (data extracted from Durand, et al. (2015), Figure 1). Starting from nearly-fixed commercial inbreds, this difference is striking. It compares to what is observed across multiple European inbreds evaluated across contrasted environments (Lehermeier, et al. 2014). Our selection experiments therefore maximized the phenotypic differences between evolved lines in nearly-identical genetic backgrounds.

**Figure 1.**
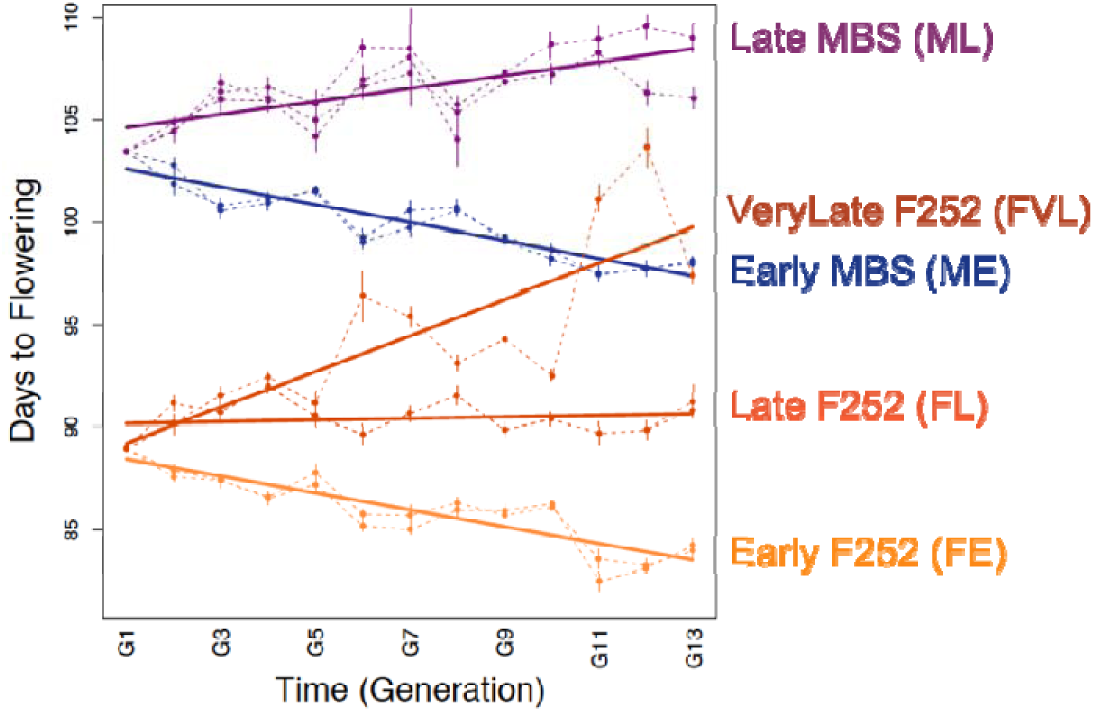
Response to selection in the Saclay divergent selection experiments during the first 13 generations. Dotted lines represent the average flowering time of each family issued from a single ancestor at G0 (Figure S1) plotted against time in generation. Flowering time was measured every year (generation) by the number of days from sowing to flowering in the experimental fields, and was corrected for the average year effect. Vertical lines indicate standard deviation within families. Data are taken from Durand et al. (2015). Plain lines represent the average trends in response to selection for the Early, Late and VeryLate F252 populations and the Early and Late MBS populations.

We chose five progenitors from the two divergent selection experiments after 13 generations of selection: one Early (FE), one Late (FL) and one VeryLate (FVL) from F252 populations, and one Early (ME) and one Late (ML) from MBS populations (Figure S1). We used offspring derived by selfing from these progenitors, first to evaluate the timing of floral transition using meristem observations and, second, to collect samples for expression analyzes. We next characterized the genome-wide transcriptomic patterns, placing a special emphasize on differentially expressed (DE) genes that have contributed to the response to divergent selection.

### Timing of floral transition

We collected from 18 to 35 shoot apical meristems from progenies of each five progenitors at different developmental stages from plants grown in the field at Université Paris-Saclay during summer 2012 and 2013 (Table 1). Plant developmental stages were defined as the number of visible leaves. We defined the shoot apical meristem Status based on shape and length as Vegetative, Transitioning or Reproductive. We determined the timing of floral transition of each progenitor as the earliest stage at which the proportion of transitioning shoot apical meristems was the highest (Figure S2, Table 1). We made three important observations from the timing of floral transition (Table 1): (1) floral transition occurred later in MBS than in F252 consistently with the flowering time difference between these two inbreds (P-value<2.2 10^-16^ and <2.35 10^-12^ in 2012 and 2013 respectively); (2) floral transition occurred at the same plant developmental stage in Early and Late genotypes in F252 (8 visible leaves), but occurred at an earlier plant developmental stage (9 visible leaves) in Early than in Late MBS progenitors (10 visible leaves); (3) the VeryLate F252 progenitor displayed a delayed timing of floral transition similar to MBS genotypes, with a year effect – floral transition occurred at 9 and 10 visible leaves for Year 1 and 2 respectively. Overall, we therefore evidenced that the selection that was applied to flowering time indirectly affected floral transition, with later occurrence of floral transition in VeryLate or Late genotypes as compared with Early.

**Table 1.**
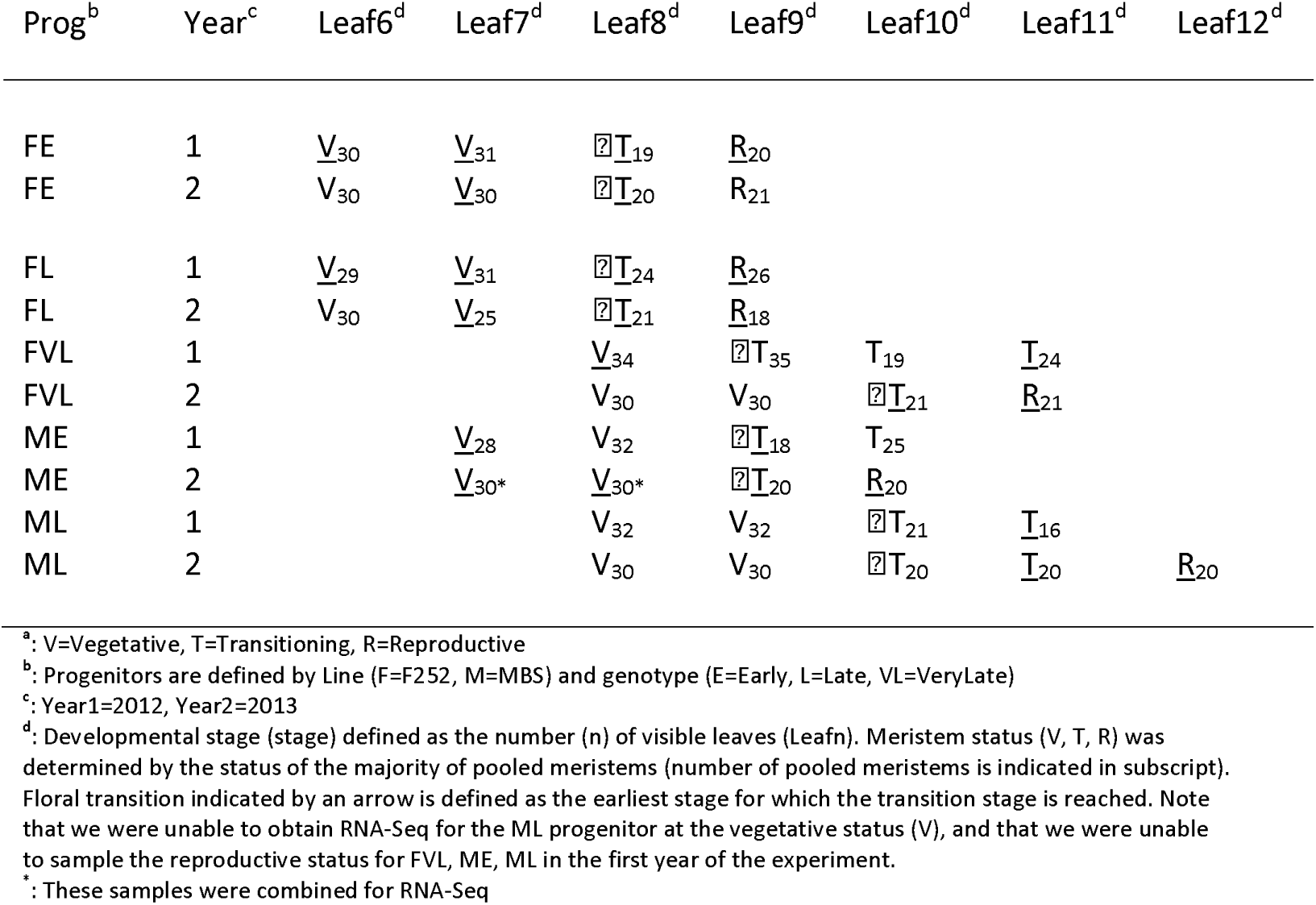
Meristem Status^a^ and timing of floral transition by Progenitor, Year and developmental stage as determined from pooled samples (samples used for RNA-Seq are underlined).

### Genome-wide patterns of gene expression in the shoot apical meristem

We pooled shoot apical meristems by developmental stage (number of visible leaves) of plants of the same progenitor to generate 25 RNA-Seq libraries (Table 1). We obtained between 19,711,727 and 33,475,403 51 bp single-reads per library (Table S1). After trimming, filtering and mapping steps, we recovered between 29.87% and 47.22% of the reads (Table S1) that were used to estimate gene expression. Hence, less than half of the original reads were used which was primarily due to “contamination” of rRNAs – around 60% of the reads corresponded to rRNAs and to a lesser extent to sequencing quality – 10% of the reads were discarded (Table S1).

We computed expression of annotated genes in the maize reference genome v3 by relying on the corresponding number of raw counts obtained from the longest transcript of each of 39,066 genes (Table S2). After filtering and normalization, we recovered a subset of 21,488 genes (55%) for which there was at least one count per million reads in half of our libraries. This estimate compares closely to the number of transcripts expressed in the maize gene atlas at two vegetative stages for pooled samples of stem and shoot apical meristem (Stelpflug, et al. 2016). We verified that the libraries considered as Replicates (same Progenitor, same shoot apical meristem Status) did cluster together, and that differences among Progenitor/Status were higher than differences among replicates (data not shown). The distributions of the normalized counts indicated lowest and highest quartiles comprised between ∼30 and ∼500, with a median depth around 200 (Figure S3). We computed pairwise correlations of normalized counts across all 25 libraries. Pearson’s correlation coefficients ranged between 0.83 and 0.99. These elevated values indicate that patterns of gene expression are overall well conserved across libraries.

### DE genes and targets of selection

We performed 27 contrasts to detect DE genes (Figure 2). In total, we detected 12,754 DE genes, 49% of which were significant in more than one contrast (Table S3). Out of the 12,754 DE genes, 3,713 displayed a Year effect, 8,228 a Line effect, and 6,568 displayed a Status effect (Figure 2). We detected 4,682 and 2,719 DE genes with a Status effect in F252 and MBS, respectively (Figure 2). Note that we detected few DE genes (<4) in contrasts between transitioning and reproductive meristems in MBS and VeryLate F252 genotypes (Figure 2). This was likely due to our inability to obtain reproductive meristems in the first year (Figure S2) that limited our power to detect DE genes between two consecutive status. The 2,499 DE genes displaying differential expression between two shoot apical meristem status in one but not all progenitors from the same line were classified in the Status x Progenitor interactions category. Finally, the Selection category regrouped DE genes exhibiting differential expression between early (FE) and late (FL or FVL) progenitors for F252, or between early (ME) and late (ML) progenitors for the MBS. Within Status x Progenitor interactions category, we also considered as part of the Selection category the subset of genes differentially expressed among Status for FE but neither for FL nor for FVL and reciprocally – DE genes among status for FL or FVL but not for FE. Such distinction was not applicable to MBS, since we detected a single DE gene between Status within ML (Figure 2). For MBS, we found 446 DE genes within the Selection category. For F252, we found 2,120 in the Selection category, that comprised 748 DE genes between E, and L or VL F252 progenitors. Considering both F252 and MBS, there were 2,451 DE genes falling into the Selection category (Figure 2 & Table S4).

**Figure 2.**
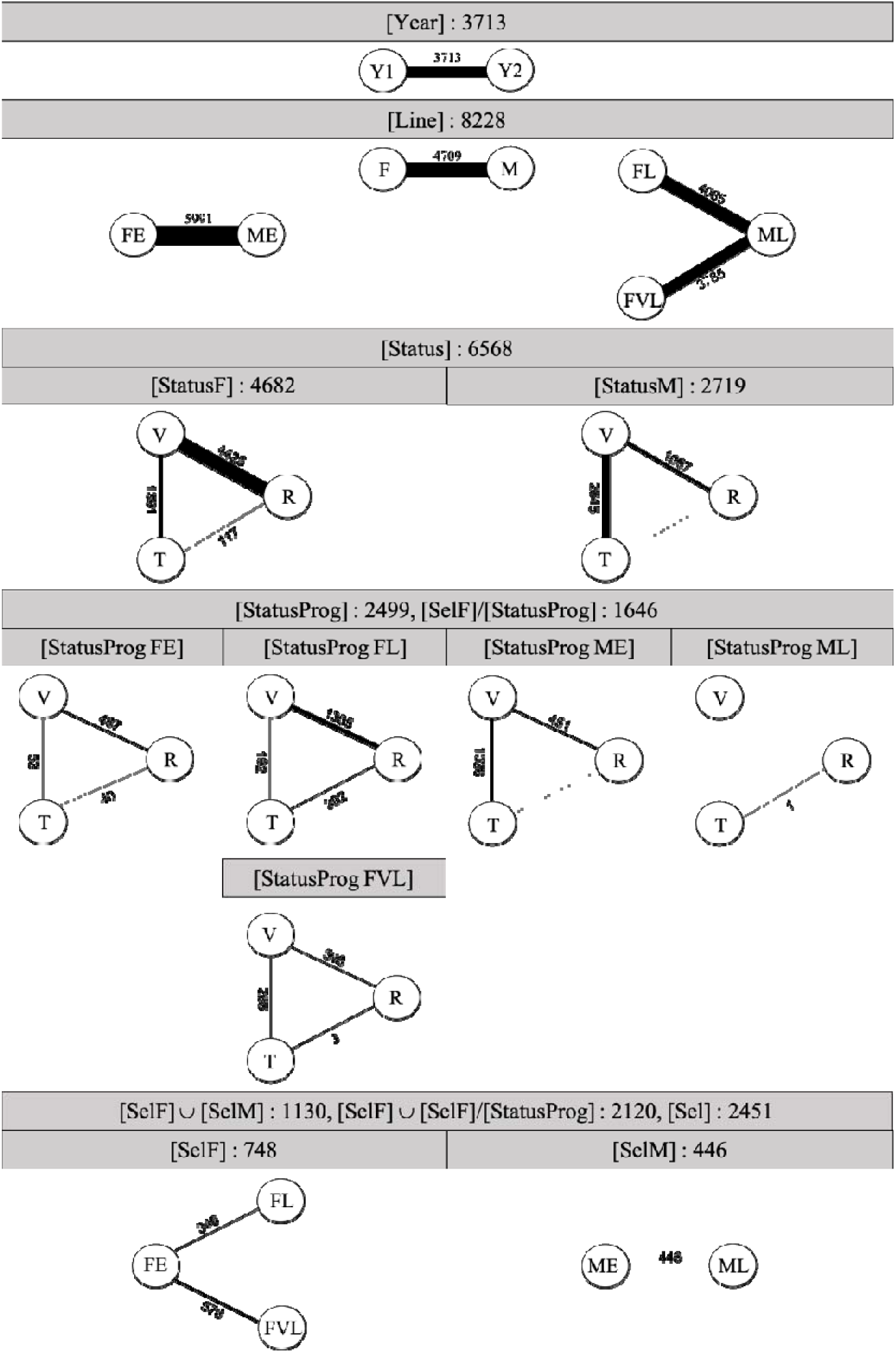
Number of Differentially Expressed (DE) genes in 27 independent contrasts. Contrasts categories and total number of detected DE genes by category (calculated from the union (⋃) of all contrasts for that category) are shown in shaded boxes. Within category, contrasted groups are connected by lines, whose thickness is proportional to the corresponding number of DE genes (indicated above/below lines with dashed line corresponding to no DE genes detected). Because we lacked the Vegetative Status for the Late Progenitor in M Line (ML), we performed a single type of contrast for [StatusProg ML]. Within the [StatusProg] category, we considered the DE genes that were differentially expressed in comparisons among status for Progenitor FE but not for Progenitors FL or FVL and reciprocally, as selected within F Line ([SelF]/[StatusProg]).

From the normalized counts, we determined for each DE gene its mean normalized expression per Progenitor per Status. We discarded from the rest of the analyses 1,481 DE genes that displayed a Year effect only, 3,262 that displayed a Line effect only as well as 641 DE genes displaying a combined Year and Line effect (Table S3 & S4). The majority (77%) of the 7,370 remaining DE genes were significant in more than one contrast. We performed a principal component analysis on the mean normalized expression of the set of 7,370 DE genes. It revealed a separation between lines and among shoot apical meristem status in F252 along axis 1, and among progenitors and status along axis 2 (Figure 3).

**Figure 3.**
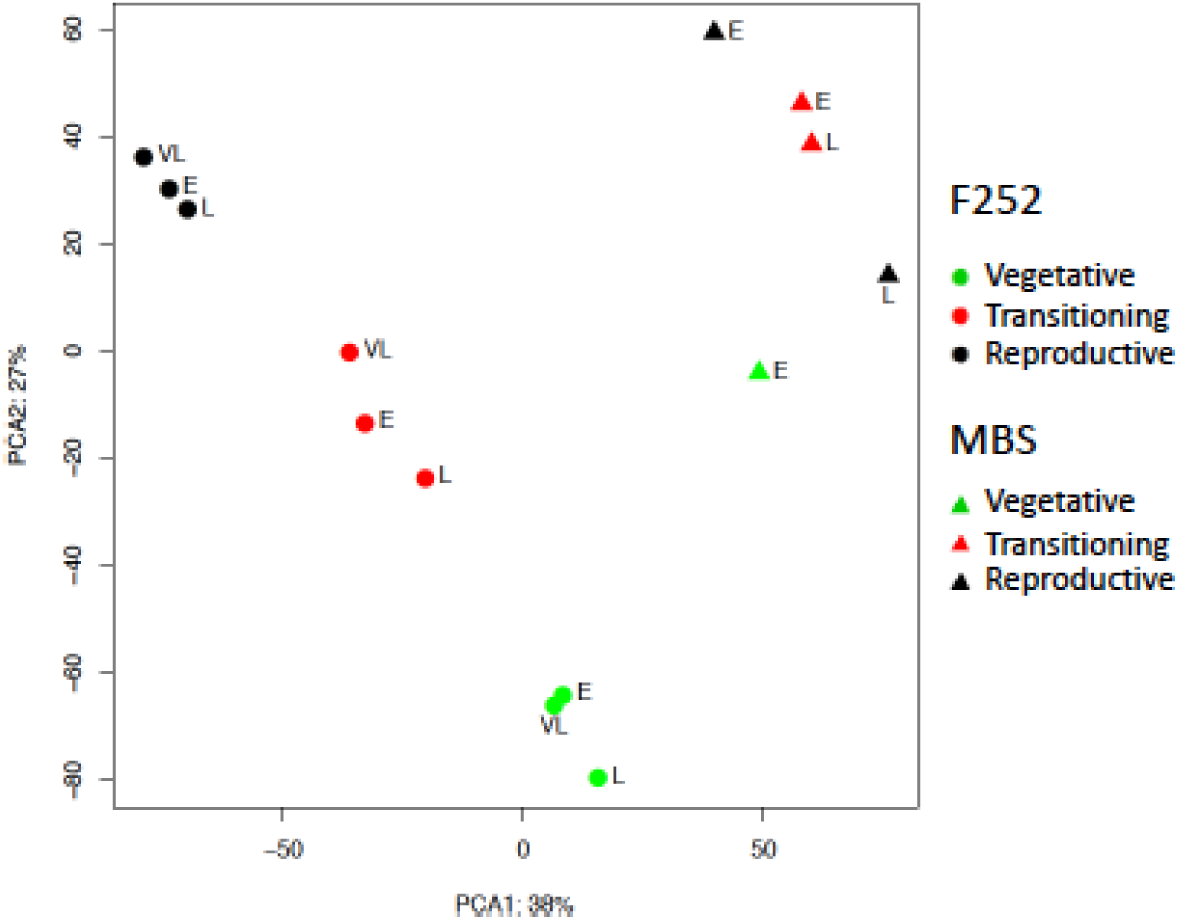
Principal component analysis on mean normalized gene expression of 7,370 differentially expressed genes, per Progenitor per Status. Values were corrected for Year effect. This subset of DE genes contains DE genes significant in at least one contrast, after discarding those displaying a Year or Line effect only, or both.

We next attributed to each gene a principal component based on correlation coefficient values (Table S4). We plotted heat maps of the 4 first components which explained respectively, 38%, 27%, 9%, 7% of the variance in gene expression. The first component (3,850 DE genes) corresponded to DE genes that differentiated lines, and shoot apical meristem status in F252 (Figure 4A). The second component (2,388 DE genes) was enriched for DE genes associated with shoot apical meristem status in F252 and in ME (Figure 4B). The third component encompassed 522 DE genes whose expression was specific to FVL progenitor as opposed to FE and FL, and varied across status in FE, FL and ME (Figure 4C). The fourth component regrouped 278 DE genes whose expression was modified during floral transition (Figure 4D). Altogether the first four components regrouped >95% of the set of 7,370 DE genes which represented 55.2% of all DE genes. Functional categories enrichment and depletion analyzes revealed specific patterns for each of the four principal components (Table S6). We found 2,451 DE genes in the Selection category. We investigated the relative contribution of genes differentially expressed between FE and FL, FE and FVL, ME and ML to the Selection category. As expected from the phenotypic response – larger between FE and FVL than between FE and FL (Figure 1) – a greater proportion of DE genes were found between FE and FVL (576) than between FE and FL (346). Comparisons of ME and ML revealed 446 DE genes (Figure 2). For F252, we found that a majority of DE genes of the Selection category were detected within the Status x Progenitor interactions category (Figure 2).

**Figure 4.**
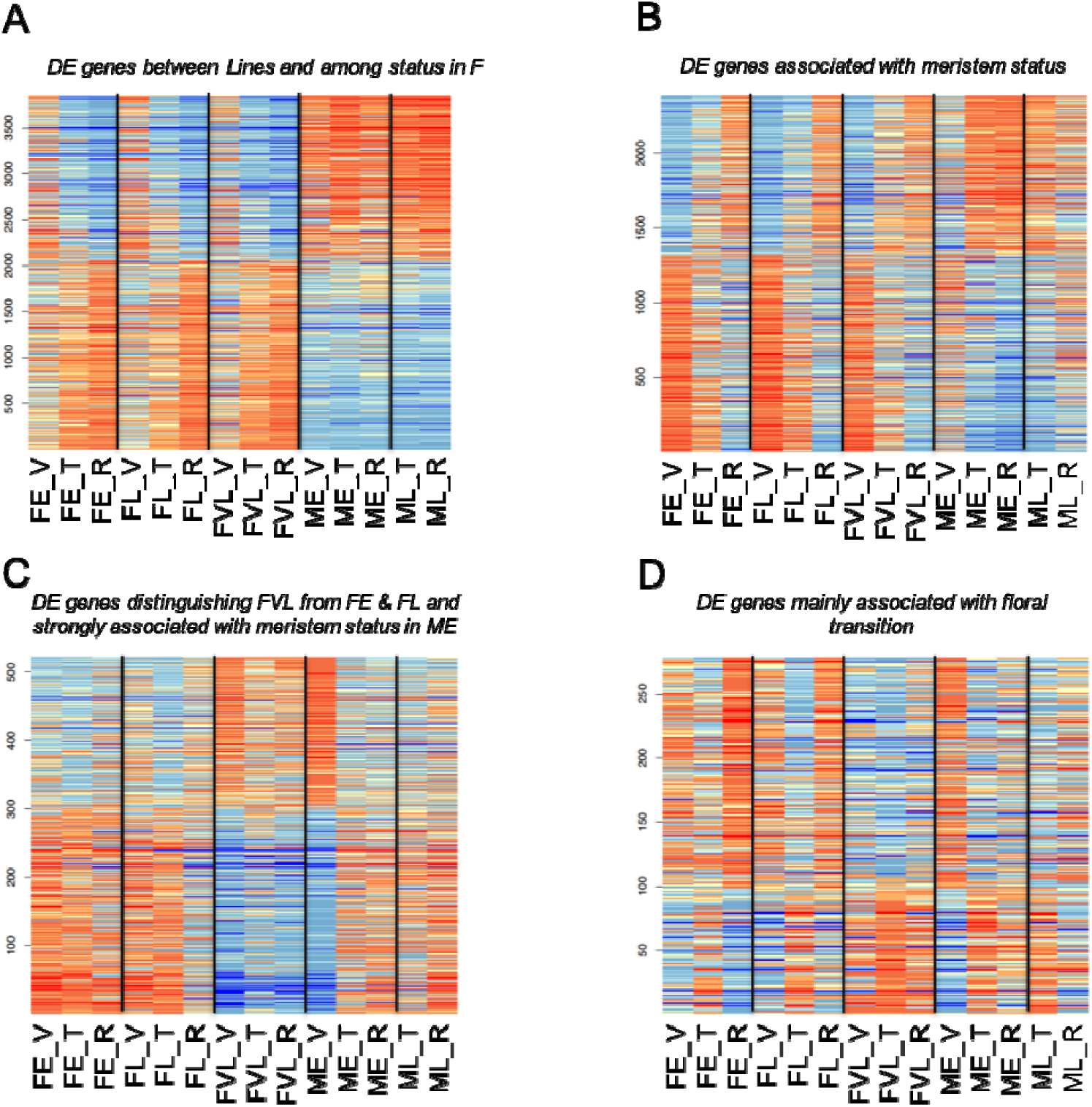
Heat maps of gene expression across Progenitors (FE, FL, FVL, ME, ML) and Status (V, T, R) for 4 independent sets of DE genes as defined by their correlations to the 4 first Principal Components. Genes are clustered by similarity of expression patterns represented here by colours (red upregulated, blue downregulated) and intensities. PC1 explained 38% of the variation (A), PC2 27% (B), PC3 9% (C) and PC4 7% (D).

To test for evidence of convergence in the response to selection between lines MBS and F252, we asked whether the count of shared DE genes belonging to the Selection category in F252 and MBS (Table S4) was significantly different from a null expectation. There were 115 shared DE genes when including Selection within Status x Progenitor interactions (for F252), and 64 shared DE genes when excluding them (Figure S4). This was significantly more than expected as tested with a two-sided exact Poisson test (with 115: expected=74 and P-value= 9.1 10^-06^, with 64: expected=26 and P-value= 3.8 10^-10^).

Altogether, our data revealed interesting features: (1) 59.3% of the 21,488 expressed genes were differentially expressed; (2) DE genes were most abundant in the Line (8,228), followed by Status (6,568) and Year (3,713) categories; (3) 2,451 genes fell in the Selection category, with MBS displaying less genes than F252; (4) There was an excess of shared DE genes in the Selection category between lines.

### Expression patterns at three genes of the gene regulatory network using qRT-PCR

In order to validate our material and methodology, we employed all RNA samples including those used to build RNAseq libraries to investigate via qRT-PCR, the expression of three genes whose interactions and effects on floral transition has been established. Those three genes are the floral meristem identity integrator *ZMM4*, the florigen *ZCN8* (Meng, et al. 2011) and the negative regulator of flowering *RAP2.7*, the two later modulating the expression of *ZMM4* (Danilevskaya, et al. 2008) (Figure S5). In addition to the shoot apical meristem, we examined expression in three additional organs: the immature part, mature part and the sheath of the last visible leaf (Figure S6).

We used *Zea mays GLYCINE-RICH PROTEIN1* (*ZmGRP1*) as a reference gene to normalize cDNA quantities. Regarding *ZmGRP1* expression in control samples (Table S6), after checking for variance and residues independence we found a significant Line (F= 242.8627, P-value= 1.249e-15) and Year effect (F= 18.3751, P-value= 0.0001828) with significant interaction (F= 191.5834, P-value= 2.621e-14) but no Plate effect (F= 0.3955, P-value= 0.5343464). Because the Plate effect was confounded with Gene effect (each gene being amplified in a different plate), the latter indicated that comparisons among our three genes were accurate. Significant differences in *ZmGRP1* expression between Lines, Years, Organs, Status:Organs (P-values<0.0428944) indicated substantial variation in the amount of cDNA quantities across samples (Table S7). We therefore normalized expression using *ZmGRP1* (corresponding ratio values (R) are provided Table S7).

Across genes, most effects (Replicate, Progenitor, Status, Organ) were significant, except for the Progenitor and the Status effects in *ZNC8* (Table S8). Patterns of normalized gene expression among replicates can be summarized as follows. The expression of *ZMM4* increased at the time of floral transition (with significant differences between vegetative and transition status) irrespective of the Organ, Progenitor and Line, and its expression was higher in the shoot apical meristem than in all other organs (Figure S7 A&D). The expression of *ZCN8* exhibited a trend towards increased expression through time in all Organs and all Progenitors. This trend was more pronounced in the shoot apical meristem between the transition and reproductive status in F252 and MBS (Figure S7 B&E). *RAP2.7* displayed a significant reduction in the shoot apical meristem at floral transition followed by a significant increase in expression at the reproductive stage. This pattern was observed in all Progenitors (Figure S7C&F) but *RAP2.7* displayed an overall lower level of expression in FVL compared with FE and FL (Figure S7C). *RAP2.7* was also more expressed in mature leaves than in any other organs.

In conclusion, qRT-PCR revealed patterns of expression consistent with *ZMM4* and *ZCN8* being floral inducers. Expression of *RAP2.7* on the other hand was repressed during floral transition. Interestingly, we noticed differences in expression between Early and Late or VeryLate Progenitors in F252 for *RAP2.7*. Note that we found good agreement between qRT-PCR and RNAseq for *ZMM4* and *RAP2.7*, with significant correlations between levels of expression determined by the two methods (Figure S8). But *ZCN8* displayed no detectable expression in RNAseq consistently with previous results detecting no expression in the meristem (Stelpflug, et al. 2016).

### Functional relevance of DE genes

In order to get more insights into the functions of detected DE genes, we undertook several approaches. First, we established from the literature a complete list of flowering time genes whose function has been at least partially validated at the molecular level (Table S9 and references herein). Note that 39 of them are already connected in the maize flowering gene regulatory network (Figure S5), and only one (GRMZM2G022489, *CORNGRASS1*) did not have raw counts information from our RNA-Seq dataset. We referred to this list of 70 genes as FT_candidates. Second, we relied on the results of the most recent GWA study on maize male and female flowering time to establish a list of 984 GWA_candidates (1001 genes were extracted from Navarro et al. (2017)), 984 of which presented raw counts information from our RNA-Seq dataset, Table S9). Third, we tested for enrichment/depletion of our DE genes in Mapman and Kegg functional categories (Table S5).

Out of 70 and 984 of the FT_candidates and GWA_candidates, 54 (representing 77.1%) and 294 (representing 29.9%) respectively displayed differential gene expression. Comparatively, FT_candidates therefore presented a greater enrichment of DE genes than GWA_candidates (P-value=7.5 10^-7^). Because the level of expression may affect our power to detect DE genes, we verified that FT_candidates were not expressed at a higher level than all transcripts taken together (P-value=0.615). After discarding the DE genes displaying either a significant Year or Line effect only, or a combination of both, there were 38 and 185 DE genes left for FT_candidates and GWA_candidates respectively. Nearly all of the 38 FT_candidates displayed differences between Status in F252 (33) and to a lesser extent in MBS (18). Remarkably, 22 of them also belonged to the Selection category in F252 (Table S9) including 6 genes connected in the maize gene regulatory network (Figure S5): *KN1, ELF4* (*EARLY FLOWERING 4 PROTEIN*), *DLF1, GIGZ1A* (*GIGANTEA 1*), *GIGZ1B* (*GIGANTEA B*), *PHYA1* (*PHYTOCHROME A1*). Likewise, for GWA-candidates, we found a majority of the remaining genes with differences between Status in F252 (119), and to a lesser extent in MBS (58) and the Selection category included 77 genes for F252, 10 genes for MBS, and two genes for both (Table S9). Comparatively, the proportion of genes falling into the Selection category among FT_candidates (22/70=31.4%) was therefore much higher than among GWA_candidates (79/984=8.0%).

## Discussion

Saclay divergent selection experiments have revealed remarkable shifts at the phenotypic level, with the production, in only 13 generations of divergent selection, of populations that flower two weeks apart in the two genetic backgrounds F252 and MBS respectively (Figure 1). Here, we showed that by selecting for flowering time difference, we indirectly selected for the timing of floral transition, which occurred earlier in early progenitors as compared with late progenitors. We used evolved lines at generation 13 to examine the transcriptomic response to divergent selection, refine our knowledge of the underlying gene regulatory network, and test whether the observed phenotypic convergence is sustained by convergence at the transcriptomic level. We generated RNA-seq from pooled samples of shoot apical meristem before/during/after floral transition. All samples were collected in the field during two consecutive years. We recovered expression for 55% of all annotated maize genes indicating that about half of them were expressed in the shoot apical meristem. We detected differential expression for roughly 59% of them.

### Expression varies primarily across developmental stages

As expected, expression varied more between lines than among developmental stages, with a proportion of DE genes exhibiting a Line or a Status effect of 64% and 51% respectively (Figure 2) confirmed by analyzes of expression patterns (Figure 3 & 4). That variation among Status concern a large proportion of genes conformed previous report (Swanson-Wagner, et al. 2012), and indicates that Status is a key component of gene expression rewiring, particularly for F252 (Figures 2 & 3).

Given the complexity of the underlying network and its regulation, we expect the expression of few genes to affect in turn the expression of a myriad of response genes. The timing of floral transition is regulated by a variety of transcriptional and post-translational regulatory mechanisms, including DNA methylation, chromatin modification, small and long noncoding RNA activity (Andres and Coupland 2012). For example, chromatin modifications in *ID1* modulate the expression of *ZCN8* in temperate maize (Mascheretti, et al. 2015). Signaling from leaf to the shoot apical meristem includes transmission of florigens (such as *ZCN8*) produced in the leaf and transported through the phloem sieve elements, making interactions with protein membranes as well as protein-protein interactions important determinants of floral transition. Finally, upregulation of protein targeting, amino acid and DNA synthesis, as well as an increase ATP production has been observed during the floral transition stage in the shoot apical meristem (Takacs, et al. 2012). As expected, we found among DE genes significant enrichment in many functional categories including amino acid metabolism, ATP synthesis, DNA, RNA, signaling, interactions, transport (Table S5).

### Reliable results obtained across two years of field experiments

While response to endogenous signals by means of the autonomous pathway (Figure S5) occupies a central role in floral transition in maize temperate lines (Colasanti and Coneva 2009), environmental signals are likely important too. We indeed found enrichment for the environmental adaptation functional category among our DE genes (Table S5). One of the main steps of maize temperate adaptation has been the rapid loss of photoperiod (Hung, et al. 2012; Teixeira, et al. 2015). Besides photoperiod, thermoregulation of flowering through the accumulation of degree days and threshold effects is also well-recognized in temperate maize and generates inter-annual variation in flowering time (Teixeira, et al. 2015). In our data, such variation influenced the timing of floral transition in the VeryLate F252 (Table 1). Surprisingly, however, this inter-annual environmental variation impacted less patterns of gene expression (29% only of all DE genes exhibited a Year effect) than developmental stages or genetic backgrounds. This important observation first indicates that RNA-seq experiments can be reliably interpreted in field conditions. Interestingly in *A. thaliana*, shoots sampled in the field at 3-days intervals over a growing season in two accessions revealed that temperature and precipitation captured a small proportion of the transcriptional variance relative to flowering status (Richards, et al. 2012). Second, this is consistent with the idea that selection during the breeding process has favored stability of expression across environments to ensure a reliable developmental outcome independently of environmental variation. Such robustness to perturbation, also called phenotypic canalization, is likely to evolve when constant phenotypic optimum – inbred line phenotype – is selected for (Abley, et al. 2016). Hence, genomic regions selected during modern breeding in temperate inbreds exhibit reduced genotype x environment interactions for grain yield, thereby limiting their plastic response (Gage, et al. 2017).

### Genes impacted by selection on flowering time are involved in floral transition transcriptional rewiring

Given our material, we paid a special attention to DE genes involved in the response to divergent selection for flowering time. Altogether they represent 19% of all DE genes, and were enriched in genes exhibiting Status x Progenitor interactions (Figure 2). Overall, F252 displayed more DE genes within the Selection category than MBS (2120 vs 446). This is in line with overall higher level of residual heterozygosity detected in the former (Durand, et al. 2015). However, if we considered only the Early and Late progenitors of each Line discarding the VeryLate Progenitor of F252, we obtained the inverse trend with less DE genes for F252 (346) than for MBS (446) (Figure 2, Figure S4). This pattern was consistent with a lack of phenotypic response in the Late F252 population after seven generations of selection but a continuation in the Late MBS population until G13 (Durand, et al. 2015). Such lack may be explained by the strong selection operated in the VeryLate F252 population that in turn relaxed selection in the Late F252 population (Figure 1).

Linkage disequilibrium is expected to be pervasive in our small populations, perpetuated by selfing. One could therefore argue that genes of the Selection category are mainly driven by fixation of large chromosomal regions during the first generations of selection generating blocks of alleles differentially expressed between Early and Late progenitors. We further tested the clustering of genes by fitting a generalized linear model where the counts of DE genes of the Selection category were determined by the number of maize genes in 1Mb non-overlapping windows and a quasiPoisson distributed error. We found an overall significant overdispersion (dispersion parameter=1.17, P-value=1.04 10^-7^), albeit with noticeable differences among chromosomes (P-value=0.0006). Chromosomes 1, 6 and 10 were significantly enriched for DE genes of the Selection category, while chromosomes 4, 7, 9 were significantly depleted.

At a first glance, these results are consistent with random fixation of segregating alleles by genetic hitchhiking accompanying the sweep to fixation of a single beneficial mutation. However, several lines of arguments support that DE genes of the Selection category do contribute to the observed phenotypic response to selection. First, the majority of these genes (76.4%) were attributed to the 2 first principal components that differentiated Status changes (Figure 4) and therefore seem to be involved in the transcriptional rewiring that occurred during floral transition. Second, these genes were depleted for “unknown” function or “not assigned” to a function (Table S5); this is consistent with an enrichment for flowering time genes that are likely well annotated. More importantly, genes of the Selection category encompassed 22 out of 54 FT_candidates that were differentially expressed, a proportion of 40.7 % that far exceeded the proportion of genes of the Selection category among all DE genes (19.2%).

### Selected DE genes from the flowering time network display patterns consistent with the literature

Among FT_candidates involved in the response to selection (Table S9), six were already connected in the gene regulatory network (Figure S5). They belong to a variety of pathways: *PHYA1* is a photoreceptor (Sheehan, et al. 2004), *KN1* belongs to the GA pathway (Bolduc and Hake 2009), *ELF4* is involved in the photoperiod pathway (Yang, et al. 2013), while *GIGZ1A, GIGZ1B* connect the latter to the circadian clock pathway (Miller, et al. 2008), and *DLF1* is a floral activator (Muszynski, et al. 2006). Flowering time is positively correlated with gibberellin accumulation in maize (Thornsberry, et al. 2001). Previous results suggest that *KN1* displays a complex pattern of expression that differs among tissues, and that it contributes to decrease gibberellin accumulation through upregulation of *GA2ox1* (Bolduc and Hake 2009). Here we found a pattern of upregulation of *KN1* expression in all progenitors during and after floral transition (Figure 5). Moreover, the expression of *GA2ox1* is downregulated during floral transition in all progenitors (Figure 5) mirroring the pattern observed in rice (Sakamoto, et al. 2001). Promotion of floral transition necessitates a critical level of *DFL1* mRNA abundance, and its expression is subsequently downregulated after (Muszynski, et al. 2006). Interestingly we found a greater level of mRNA abundance before floral transition in VeryLate progenitor compared with the other progenitors, suggesting that the former has evolved a distinct threshold level above which floral transition is initiated.

**Figure 5.**
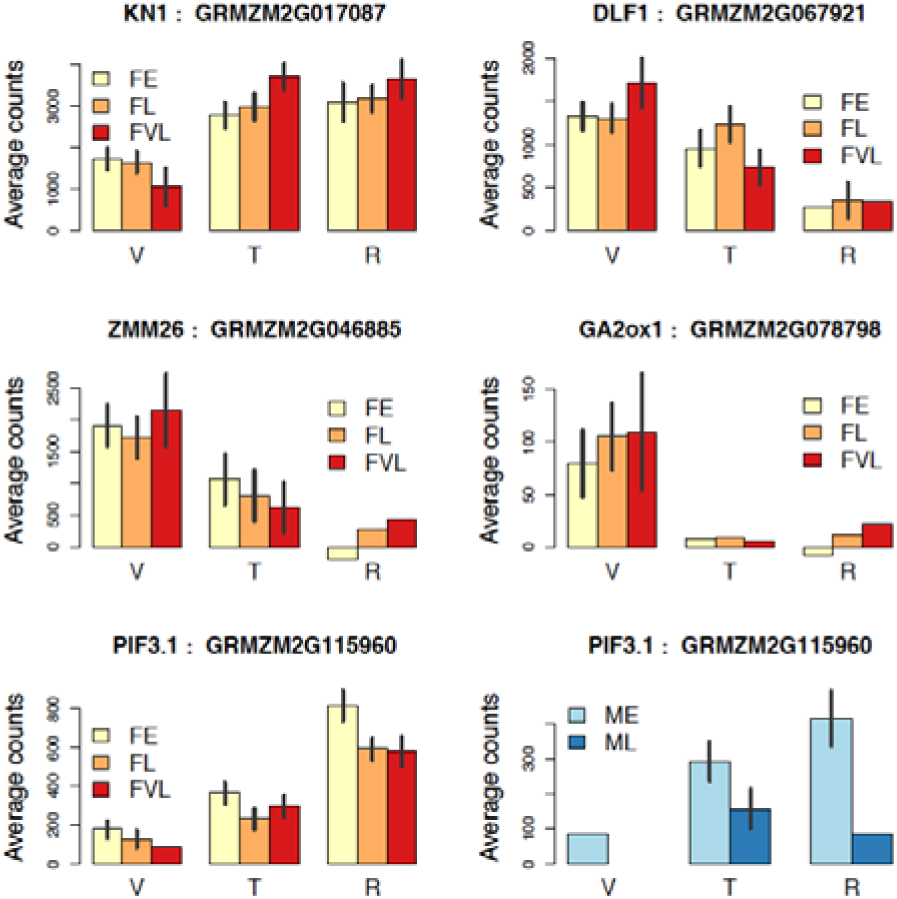
Expression patterns of 5 maize flowering time genes and candidates as determined by RNA-seq. Expression was determined by shoot apical meristem Status (V=Vegetative, T=Transitioning, R=Reproductive) and Progenitor in the two Divergent Selection Experiments (F252 and MBS in red and blue colors, respectively) for *KN1, GA2ox1, DLF1, ZMM26, PIF3.1*. Expression is measured by average counts across replicates (after correcting for the year effect) and vertical bars represent the standard deviation around the mean, obtained after dividing the residual variance estimated by DESeq2 for each gene by the number of replicates.

The overall observed patterns were consistent with the roles of *ZMM4* and *ZCN8* in promoting floral transition in temperate maize, with upregulation of *ZMM4* in all organs at floral transition – including the shoot apical meristem – in both F252 and MBS (Danilevskaya, et al. 2008) (Figure S7A&D), and increased expression of *ZCN8* during floral transition (Meng, et al. 2011) in the shoot apical meristem also in both inbred lines (Figure S7B&E). In addition, we report here the first qRT-PCR survey of the expression of *RAP2.7* in the shoot apical meristem during floral transition (Figure S7C&F). It displayed patterns consistent with its role as a negative regulator of flowering time (Salvi, et al. 2007). Interestingly, our qRT-PCR results suggested a potential role of *RAP2.7* in the response to selection. We were unable to corroborate this result using RNAseq because the level of expression of *RAP2.7* was very low (median of 3 counts across libraries Table S2, Figure S8). Consistently, a previous study detected very low/no expression (fragments per kilobase exon model per million mapped fragments [FPKM]<2) using RNAseq from meristem and leave tissues (Stelpflug, et al. 2016). Previous studies have shown that the presence of miniature transposon in a conserved non-coding sequence upstream *RAP2.7* is associated with early-flowering (Salvi, et al. 2007; Ducrocq, et al. 2008). More recently a model has been proposed whereby methylation spreading from the transposon reduces the expression of the *RAP2.7* in the leaves causing early flowering (Castelletti, et al. 2014). In contrast, we found a lower expression level in VeryLate F252 than in Late F252 in all organs. Because the MITE insertion is present in F252 (Ducrocq, et al. 2008) and unlikely polymorphic among our samples, differential expression among alleles may be caused by a complex regulation including the action of miR172 (Figure S5).

Besides those FT_candidates already connected in the gene regulatory network, others merit attention. For instance, we detected the three *RAMOSA* genes (Table S9) that act together to determine inflorescence branching (Vollbrecht, et al. 2005; Bortiri, et al. 2006), one of which, *RA1*, may be in part regulated by sugar signaling (Satoh-Nagasawa, et al. 2006). In addition, we confirmed the suspected role of *ZMM26* (GRZM2G046885) in maize flowering time transition (Alter, et al. 2016) as it was found under selection in F252 (Table S9). The more pronounced downregulation of *ZMM26* at the time of floral transition may be involved in delayed flowering in the VeryLate progenitor (Figure 5). Finally, PIF3.1, a member of a family of phytochrome interacting factor in maize that mediates plant response to various environmental factors (Kumar, et al. 2016), was found under selection in F252. The trend towards increased expression in the shoot apical meristem at floral transition in the Early progenitors of both inbreds indicates a potential role in promoting early flowering (Figure 5).

### Evidence of genetic convergence between inbred lines

We detected convergence of transcriptome response between lines submitted to the same selection pressure, which translated into an excess of shared DE genes of the Selection category between lines (115 genes). Such convergence of molecular phenotypes may result from distinct regulatory mechanisms of gene expression: selection of mutations in trans-acting factors that modulate expression of entire pathways (Li, et al. 2013); or selection of mutations in cis-acting factors. An example of the latter was recently illustrated by the selection of two independent cis-regulatory variants at the flowering time gene *ZCN8* during early domestication and later diffusion of maize (Guo, et al. 2018). Interestingly, these two variants pre-existed before the onset of selection in teosinte populations.

Likewise, in our Divergent Selection Experiments, we found an example of selection in a genomic region encompassing several genes among which *EiF4A*, which displayed residual heterozygosity in the initial F252 seed lot. This region subsequently underwent differential fixation of the two alleles in the Early and VeryLate F252 progenitors (Durand, et al. 2012), and we actually found evidence for differential gene expression of *EiF4A* between Early and Late/VeryLate progenitors (Table S5). Note that in maize, stretches of residual heterozygosity have been shown to be either unique or shared by very few lines (Brandenburg, et al. 2017). Therefore, except for shared streches between F252 and MBS, sorting of pre-existing alleles by differential selection between early and late populations should not translate into patterns of convergence between inbred lines.

While convergence between inbred lines can happen from shared standing genetic variants, previous results have shown that *de novo* mutations have contributed to our observed response to selection. Convergence of gene expression between our populations may therefore also originate from such mutations acquired independently across lineages. Examples of such convergence exist *in natura* such as the well-known case of pelvic reduction in freshwater stickleback (Chan, et al. 2012). In experimental evolution systems such as yeast or bacteria, parallel *de novo* mutations have been often observed at the gene and even nucleotide level during adaptation to antibiotic dosage (Laehnemann, et al. 2014), to nutrient availability and growth conditions (Spor, et al. 2014; Turner, et al. 2018), and during acclimation to high temperature (Tenaillon, et al. 2012). In this last example, first-step mutations in the evolved lines have been observed in multiple replicates at a single gene: a transacting factor modulating the expression of hundreds of downstream genes (Rodriguez-Verdugo, et al. 2016). Such pattern is consistent with our observations with convergence of expression at both few candidates and many response genes.

The relative contribution of standing expression variants in ancestral genotypes versus *de novo* expression variants acquired independently during the course of the experiment, to patterns of convergence remains to be established. This could be achieved by investigating in the ancestor lines, allele specific expression of the subset of 115 DE genes that displayed convergence of expression between the two inbreds. In addition, allele specific expression in hybrids created from crosses between evolved genotypes would bring insights into cis-versus trans-regulation of gene expression (de Meaux, et al. 2005); and whole genome sequencing of ancestral and derived genotypes would allow identifying the origin of mutations and their fate through generations of selection.

## Conclusion

We used two divergent selection experiments with controlled genetic backgrounds and little residual heterozygosity to characterize the genome-wide transcriptomic response to selection for flowering time. Throughout this study, we have demonstrated the reliability of performing RNA-seq experiments in field conditions. We have shown that the meristem developmental status is the second main source of differential gene expression. We uncovered a subset of genes involved in the response to selection. This subset of genes likely encompasses a majority of response genes; but also displays a strong enrichment for flowering time genes with evidence of convergence of expression between lines sustaining phenotypic divergence. Modeling of the floral transition gene network would help dissociating causal from response genes.

## Material and Methods

### Plant Material

We have conducted two independent divergent selection experiments for flowering time from two commercial maize inbred lines, F252 and MBS847 (MBS). These experiments were held in the field at Université Paris-Saclay (Gif-sur-Yvette, France). Within each divergent selection experiments, plants selected as early- or late-flowering were selfed at each generation, and offspring used for the next generation of selection. As a result, we derived from each divergent selection experiment, two populations of Early- and Late-flowering genotypes previously identified as Early F252, Late F252, Early MBS, Late MBS (Durand, et al. 2015), as well as a VeryLate F252 populations. Seeds from selected genotypes at all generations were stored in cold chamber. We traced back the F252 and MBS pedigrees from generation 13 (G13) to the start of the divergent selection experiments (Durand, et al. 2015, Figure S1).

In order to investigate genome-wide changes of gene expression before, during, after flowering transition and contrast those changes between Early and Late genotypes, we chose five progenitors selected at G13 from the two divergent selection experiments: one progenitor from the Early (FE), Late (FL) and Very Late (FVL) F252 populations, and from the Early (ME) and Late (ML) MBS populations (Figure S1).

All 5 progenitors were selfed to produce seeds. The resulting progenies were sown and grown in the field at Université Paris-Saclay (Gif-sur-Yvette, France) during summer 2012 and 2013. The experimental design contained rows of 25 plants from the same progenitor. For each line, each of the progenitors was represented by nine rows that were randomized in three blocks. Precipitations over the growing period totalized 117 mm and 60.5 mm and the average daily temperature reached 16.51 °C and 18.41 °C for 2012 and 2013, respectively. The second year of experiment was therefore hotter and drier than the first year. We added as controls, plants from F252 and MBS initial seed lots.

We defined developmental stages as the number (n) of visible leaves — the cotyledon leaf being considered as the first one, and named them accordingly Leafn. Based on preliminary observations, we defined 4 to 5 developmental stages per progenitor from leaf6 to leaf12 that encompassed the flowering transition (Table 1). Between June 08^th^ and July 9^th^ for 2012 (Year 1) and June 14^th^ and July 19^th^ for 2013 (Year 2), we collected plants from the different blocks on a daily basis early morning (between 8:00 and 9:00 am). We randomly chose plants at the same developmental stage for a given progenitor. We dissected four organs from the fresh material collected: the shoot apical meristem, the immature and mature part of the last visible leaf (IL, ML), the sheath at the basis of the last visible leaf (S). We recorded the developmental stage (number of visible leaves) as well as the shoot apical meristem status: Vegetative (V), Transitioning (T) or Reproductive (R). We established shoot apical meristem status based on its shape following (Irish and Nelson, 1991) and length. Typical range of lengths are V: 0.1-0.2 mm, T: 0.2-0.35 mm, R>0.35 mm.

For a given organ, we pooled together from 18 to 35 plants from the same progenitor at the same developmental stage (Table 1). We used as controls, plants from the original seed lots for F252 and MBS. These controls were dissected only at stage leaf9 and leaf11 respectively, and all four organs were pooled together within a tube. All collected organs were frozen in liquid nitrogen upon collection and stored at −80 °C. In total, we gathered five progenitors x 4 organs x 4 stages that are 80 samples per year in addition to the F252 and MBS controls. In 2013, we included one more stage for the ML progenitor for a total of 86 samples including the two controls. We performed a single biological replicate in 2012 (Replicate 1) and two biological replicates in 2013 (Replicates 2 and 3) for all organs, except the shoot apical meristem for which a single biological replicate was done each year. Total RNAs of pooled samples were extracted using Qiagen RNeasy Plant kit for shoot apical meristems, and using TRIzol reagent (Invitrogen) and ethanol precipitation for other organs. Total RNA was treated with DNAse (Ambion) following the manufacturer’s instructions. We evaluated RNA yields using Nanodrop 2000 (Thermo Scientific).

### qRT-PCR assays and statistical procedure

For all samples, 4 µg or less (for samples with <4 µg) of total RNA was reverse transcribed using random hexamers (Thermo scientific), 200 units of Revertaid Reverse Transcriptase (Thermo Scientific), and 20 units of recombinant Rnasin RNase inhibitor (Promega) in a final volume of 20 µl. Simultaneously, all samples were subjected to the same reaction without the RT to verify that there was no genomic DNA contamination. We designed copy-specific primers to amplify the following 5 genes: *MADS-transcription factor 4* (*ZMM4*, GRMZM2G032339), *phosphatidylethanolamine-binding protein8* (*ZCN8*, GRMZM2G179264), *Apetala-2 domain transcription factor* (*RAP2.7*, GRMZM2G700665). Primer sequences are the following *ZMM4* F : 5’GGAGAGGGAGAAGGCGGCG 3’, R : 5’ CTACTCAAGAAGGCGCACGA 3’; *ZCN8* F : 5’ATGCGCCACAACTTCAACTG 3’, R : 5’GAAGAGTAGAAACCATAGGCCACTGA; 3’*RAP2.7* F : 5’CGCCGACATCAACTTCAACC 3’, R : 5’CTCCAGGTACAGAGGCGTCA 3’. We used *Glycine-rich protein1* (*ZmGPR1=GRMZM2G080603*) with the following primers F:5’-CACAACGCCTTCAGCACCTA-3’, R:5’-AAGGTGACGAAGCCGAAGC-3’ as a reference gene with ubiquitous expression among stages and organs following (Virlouvet, et al. 2011). *ZmGRP1* has been shown to be the second most stably expressed genes among 60 distinct tissues representing different organs x stages (Sekhon, et al. 2011).

We used standard qRT-PCR protocols with the SYBR Green PCR Master Mix (Applied biosystems) and the 7500 Real Time PCR System (Applied Biosystems) to evaluate gene expression. We undertook calibration procedure to ensure equivalent PCR efficiency across genes using serial 7-fold dilutions of cDNAs. We verified the specificity of the amplification by dissociation curve analysis, gel electrophoresis and sequencing of PCR products. We used a single 96 deep-well plate per gene to PCR amplify samples of a given replicate, the F252 and MBS controls, one negative control (without cDNA) to verify that there was no genomic DNA contamination. In addition, we amplified the F252 and MBS control with the *ZmGRP1* reference gene in every plate.

We quantified gene expression by the amplification Cycle threshold (C_T_). C_T_ indicates the number of cycles at which Q_0_=Q_CT_/(1+*E*)^CT^. We first used *ZmGRP1* C_T_ in F252 and MBS controls to evaluate the significance of the effects of the Line (F252, MBS), Year (1, 2) and their interaction, as well as the “deep-well plate effect” (Plate 1-6) confounded with the Gene effect. Second, we used *ZmGRP1* C_T_ of all samples (except controls) to test significance of the Replicate, Progenitor, shoot apical meristem Status (V, T, R), Organ (shoot apical meristem, Immature Leaf, Mature Leaf and Sheath) effects and the interactions between Status:Organ. In order to account for differences in initial cDNA quantities across samples, we normalized C_T_ across all samples by *ZmGRP1* C_T_ values. Resulting normalized values (Ratios=R) were used in all subsequent analyses.

For all three genes, we employed the following linear regression model that decomposed variations of R into 4 fixed effects, an interaction effect and a residual:

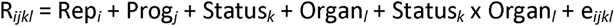

with Rep_*i*_ =1,2,3; Prog_*j*_ = FE, FL, FVL, ME, ML; Status_*k*_ =V, T, R; Organ_*l*_ =M, IL, ML, S; and e_*ijkl*_ the residual. We calculated adjusted means after correcting for Rep effect, and 95% confidence intervals. Finally, we performed contrasts to compare adjusted means of all 60 pairwise combinations of Progenitor (5) × Status (3) × Organ (4). We retained as significant P-values<1/1000 which was just above the Bonferroni threshold.

### RNA sequencing, data filtering and mapping

We used a subset of the RNA samples used in the qRT-PCR to perform RNA-Seq. This subset contained shoot apical meristem samples from 2012 and one of the 2013 replicate. Because we were limited in the amount (after qRT-PCR reactions) and quality of total extracted RNAs, we were able to obtain sequencing data for a subset of 25 samples (25 libraries, Table 1). Note that we combined Leaf7 and Leaf8 for ME in 2013 in a single sample. In brief from 2.5 to 5 µg of total RNA, mRNA was isolated with two rounds of polyA purification, fragmented, converted to cDNA, and PCR amplified according to the Illumina RNA-Seq protocol (Illumina, Inc. San Diego, CA). Oriented cDNA libraries were constructed and 51 bp single-reads sequences were generated using the Illumina Genome Analyzer II (San Diego, CA) at the high throughput sequencing platform of IMAGIF (Gif-sur-Yvette). Illumina barcodes were used to multiplex the samples.

From raw reads, we retrieved Illumina adapters using the software Cutadapt 1.2.1 (Martin 2011) with the following parameters: *count=5* ; *minimum length=10* ; *length overlap=3*. We trimmed low quality reads by imposing 3 consecutive bases at the 3’ end with a score of >20 and discarding reads with a sequence length after trimming <25 pb. We filtered reads with a perfect match against maize rRNAs. We mapped the resulting reads (trimmed, filtered and uniquely-mapped) against the maize reference genome v3 (http://ftp.maizesequence.org/) using Tophat (Trapnell, et al. 2009) that accounts for exons-introns junctions using end-to-end alignment with the following parameters adapted to shorts reads with a short seed (L15) and with very sensitive parameters (-i): *--b2-D 25 --b2-R 3 --b2-N 0 --b2-L 15 --b2-i S,1,0.5 --b2-n-ceil L,0,0.5.* We provided transcript annotations during the reads mapping. From the resulting sam files we retrieved the uniquely mapped reads imposing a filter on read containing NH:i:1. We used coverageBed with the –split option where bed entries are treated as distinct intervals, and retained reads mapped to exons only (as described in Zea_mays.AGPv3.22.gtf, http://ftp.maizesequence.org/). We parsed the resulting file to determine the number of exons, the transcript length, the number of counts (raw counts) per transcript and their coverage (number of bases covered by at least one read). We used the raw counts of the longest transcript of each gene to perform the rest of the analyses.

### RNA-Seq filtering and normalization

We used the standard procedures proposed by DESeq2 v1.14.1 (Love, et al. 2014) for filtering and normalization. We computed libraries size factors after data filtering to eliminate transcripts with less than one count per million reads in half of the libraries. We considered libraries of the same progenitor having the same shoot apical meristem status albeit different leaf stages as replicates irrespective of the year of experimentation. We applied the following graphical procedures to verify our pipeline: we examined boxplots of the libraries counts distributions to validate the normalization procedure; we evaluated pairwise correlations between replicates both on normal and logarithmic scales; we performed multidimensional scaling (MDS) as suggested to compute pairwise distances between libraries based on the 500 genes with the largest standard deviation among samples, and represented the libraries graphically in a two dimensional space.

### Detection of Differentially Expressed genes (DE genes) from RNA-Seq data

In order to identify Differential Expressed genes (thereafter, DE genes), we used the DESeq2 package (Love, et al. 2014) along with its internal filtering procedure that chooses the filtering threshold from the mean gene expression of DE genes. The overdispersion parameters were estimated from the whole data set. We next performed pairwise comparisons between libraries by computing contrasts sequentially and determining the associated P-values for each transcript after correction for multiple testing (Benjamini and Hochberg 1995).

In our experimental setting, a Progenitor represents an early or a late genotype (E, L, or VL) issued from one of the two Lines (F or M). In addition, we sampled three shoot apical meristem Status (V, T, R) and performed two Years of field experimentation. Altogether, we tested the following contrasts: (1) one comparison between Year effect averaged across all samples; (2) one comparison between Line effect averaged across all samples ; (3) three independent comparisons between early and late Progenitors of each Divergent Selection Experiment regardless of the shoot apical meristem Status (FE vs FL, FE vs FVL, ME vs ML); (3) three independent comparisons between Lines regardless of the shoot apical meristem Status considering either early and late Progenitors alternatively (FE vs ME, FL vs ML, FVL vs ML); (4) three independent comparisons of shoot apical meristem Status within each Divergent Selection Experiment regardless of Progenitors (V vs T, V vs R, T vs R); (5) three independent comparisons of shoot apical meristem Status by Progenitor (V vs T, V vs R, T vs R in FE, FL, FVL, ME, ML, respectively). Considering 25 samples and missing combinations of Progenitor × Status × Year (Table 1), we performed a total of 27 independent contrasts. Note that the filtering procedure generated different sets of genes retained for each contrast.

Altogether, our analysis corresponded to the following linear decomposition of the mean expression, *Theta*_*ylgs*_:

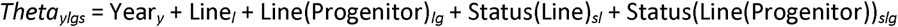

We determined the number of Differentially Expressed (DE) genes in each contrast. We next considered as DE genes the ones that were detected as differentially expressed in at least one contrast. From the normalized counts, we determined for each DE gene its mean expression per Progenitor per Status, corrected for the Year effect using a linear model. We further normalized mean gene expression by the overall mean, so that a negative value corresponds to under-expression, and a positive value to over-expression.

### Gene clustering from differential expression patterns

We performed a Principal Component analysis (PCA) using mean normalized gene expression of DE genes, after discarding those displaying a Year or Line effect only, or both. Principal components were defined as linear combinations of genes and allowed projection of Progenitor × Status combinations. To reduce the complexity, we used an approach similar to the one proposed in the MixOmics package (Rohart, et al. 2017). We calculated the Pearson correlation coefficient of each gene to the first 13 Principal Components and, based on the absolute value of the greatest correlation coefficient among 13, we attributed each gene to a single principal component axis. Correlation coefficients were preferred to PCA loadings because they accounted for both the variance of the PC axis and the trait variance. We retained 4 lists of DE genes corresponding to the 4 first principal components and built independent heat maps after ordering genes according to their correlation coefficient – from positive to negative correlations.

### Functional analysis of DE genes

We conducted a functional annotation of DE genes using the MapMan (http://mapman.gabipd.org/web/guest/mapman) and Kegg Ontology level 1, 2 and 3 (https://www.genome.jp/kegg/ko.html). We reduced the number of functional categories by reassigning each of the 226 Mapman categories to 36 Mapman categories, and each Kegg levels 2 and 3 to 24 Kegg pathways, corresponding to the following five high-level kegg categories (Kegg level 1): Metabolism, Genetic Information processing, Environmental Information processing, Cellular processes, Organismal systems (Table S10). We next used a bilateral exact Fisher to test for categories’ enrichment/depletion for DE genes belonging to the 4 first principal components, DE genes within the Selection category and genes differentially expressed between lines most of which were discarded from the PCA analysis. We computed the standardized residuals as a standardized measure of the difference between the observed and expected counts.

## Data accessibility

RNA-seq data were deposited in the Gene Expression Omnibus database from the National Center for Biotechnology Information under SRA accession number: PRJNA531088.

## Supplementary material

### Supplementary tables and R script are available at

https://figshare.com/articles/Supplementary_Tables/7271399/4 (DOI:10.6084/m9.figshare.7271399)

Table S1. Sequencing and mapping statistics.

Table S2. Raw counts for the longest transcript of 39,066 gene in all 25 libraries.

Table S3. Number of Differentially Expressed (DE) genes in 27 contrasts.

Table S4. List of Differentially Expressed genes (12,754) with average gene expression and standard deviation.

Table S5. Enrichment tests of Mapman categories and Kegg pathways for DE genes.

Table S6: Expression of the reference gene *ZmGRP1* as C_T_ in F252 and MBS controls.

Table S7: Expression of the reference gene *ZmGRP1* and 5 candidate genes as measured by CT in all samples, and normalized expression in candidates.

Table S8: Significance of effects for each of the 5 candidate genes as determined by linear regression.

Table S9. List of 71 flowering time genes in maize and overlap with Differentially Expressed genes. (continued) List of 2001 Female and/or Male Flowering Time candidates in maize and overlap with Differentially Expressed genes.

Table S10. Functional annotations of maize transcripts based on Mapman and Kegg ontologies.

### Supplementary figures are available in the Supplemental material

Figure S1. Pedigree of the progenitors in the F252 and MBS divergent selection experiments.

Figure S2. Proportion of Status among all dissected meristems (Table 1) in pooled samples for Year 1 and Year 2 by Progenitor FE (A), FL (B), FVL (C), ME (D), ML (E).

Figure S3. Distribution of normalized counts with median values (dots), 25 and 75 quantiles (vertical lines), for the 25 RNA-seq libraries.

Figure S4. Venn Diagram of DE genes included in the Selection category.

Figure S5. Schematic representation of maize flowering time pathway.

Figure S6. Organs used for the qRT-PCR in addition to the shoot apical meristem.

Figure S7. Adjusted means of log(Expression) across Replicates and 95% CI determined by qRT-PCR for 3 genes *ZMM4, ZCN8, RAP2.7* in F252 (A-C) and MBS (D-F).

Figure S8. Correlations between levels of expression determined by qRT-PCR and RNA-seq for two candidate genes.

## Acknowledgements

This work has benefited from the facilities and expertise of the high throughput sequencing platform of IMAGIF (Centre de Recherche de Gif, www.imagif.cnrs.fr), with appreciated advice from Delphine Naquin, Maud Sylvain and Yan Jaszczyszyn. We are grateful to Coraline Linguat, Betty Leitte, and Hélène Corti for their technical assistance; and to Peter Morrell for helpful discussion. We would like to thank Tanja Pyhäjärvi, Laura Shannon and two anonymous reviewers for very insightful reviews and careful editing of our manuscript. This preprint has been peer-reviewed and recommended by Peer Community In Evolutionary Biology (https://doi.org/10.24072/pci.evolbiol.100071)

## References

Abley K, Locke JCW, Leyser HMO. 2016. Developmental mechanisms underlying variable, invariant and plastic phenotypes. Annals of Botany 117:733–748.

Alter P, Bircheneder S, Zhou L-Z, Schlueter U, Gahrtz M, Sonnewald U, Dresselhaus T. 2016. Flowering Time-Regulated Genes in Maize Include the Transcription Factor ZmMADS1. Plant Physiology 172:389–404.

Andres F, Coupland G. 2012. The genetic basis of flowering responses to seasonal cues. Nature Reviews Genetics 13:627–639.

Benjamini Y, Hochberg Y. 1995. Controllling the false discovery rate - a practical and pwerful approach to multiple testing. Journal of the Royal Statistical Society Series B-Methodological 57:289–300.

Bolduc N, Hake S. 2009. The Maize Transcription Factor KNOTTED1 Directly Regulates the Gibberellin Catabolism Gene ga2ox1. Plant Cell 21:1647–1658.

Bortiri E, Chuck G, Vollbrecht E, Rocheford T, Martienssen R, Hake S. 2006. ramosa2 encodes a LATERAL ORGAN BOUNDARY domain protein that determines the fate of stem cells in branch meristems of maize. Plant Cell 18:574–585.

Bouche F, Lobet G, Tocquin P, Perilleux C. 2016. FLOR-ID: an interactive database of flowering-time gene networks in Arabidopsis thaliana. Nucleic Acids Research 44:D1167–D1171.

Brandenburg J-T, Mary-Huard T, Rigaill G, Hearne SJ, Corti H, Joets J, Vitte C, Charcosset A, Nicolas SD, Tenaillon MI. 2017. Independent introductions and admixtures have contributed to adaptation of European maize and its American counterparts. Plos Genetics 13(3):e1006666.

Buckler ES, Holland JB, Bradbury PJ, Acharya CB, Brown PJ, Browne C, Ersoz E, Flint-Garcia S, Garcia A, Glaubitz JC, et al. 2009. The Genetic Architecture of Maize Flowering Time. Science 325:714–718.

Burke MK, Dunham JP, Shahrestani P, Thornton KR, Rose MR, Long AD. 2010. Genome-wide analysis of a long-term evolution experiment with Drosophila. Nature 467:587–590.

Burke MK, Liti G, Long AD. 2014. Standing Genetic Variation Drives Repeatable Experimental Evolution in Outcrossing Populations of Saccharomyces cerevisiae. Molecular Biology and Evolution 31:3228–3239.

Castelletti S, Tuberosa R, Pindo M, Salvi S. 2014. A MITE Transposon Insertion Is Associated with Differential Methylation at the Maize Flowering Time QTL Vgt1. G3-Genes Genomes Genetics 4:805–812.

Chan YF, Jones FC, McConnell E, Bryk J, Buenger L, Tautz D. 2012. Parallel Selection Mapping Using Artificially Selected Mice Reveals Body Weight Control Loci. Current Biology 22:794–800.

Chan YF, Marks ME, Jones FC, Villarreal G, Jr., Shapiro MD, Brady SD, Southwick AM, Absher DM, Grimwood J, Schmutz J, et al. 2010. Adaptive evolution of pelvic reduction in sticklebacks by recurrent deletion of a Pitx1 enhancer. Science 327:302–305.

Colasanti J, Coneva V. 2009. Mechanisms of Floral Induction in Grasses: Something Borrowed, Something New. Plant Physiology 149:56–62.

Colasanti J, Yuan Z, Sundaresan V. 1998. The indeterminate gene encodes a zinc finger protein and regulates a leaf-generated signal required for the transition to flowering in maize. Cell 93:593–603.

Coneva V, Guevara D, Rothstein SJ, Colasanti J. 2012. Transcript and metabolite signature of maize source leaves suggests a link between transitory starch to sucrose balance and the autonomous floral transition. Journal of Experimental Botany 63:5079–5092.

Danilevskaya ON, Meng X, Selinger DA, Deschamps S, Hermon P, Vansant G, Gupta R, Ananiev EV, Muszynski MG. 2008. Involvement of the MADS-Box gene ZMM4 in floral induction and inflorescence development in maize. Plant Physiology 147:2054–2069.

de Meaux J, Goebel U, Mitchell-Olds T. 2005. Allele-specific assay reveals functional variation in the Chalcone Synthase promoter of Arabidopsis thaliana that is compatible with neutral evolution. Plant Cell 17(3):676–690.

Dong Z, Danilevskaya O, Abadie T, Messina C, Coles N, Cooper M. 2012. A gene regulatory network model for floral transition of the shoot apex in maize and its dynamic modeling. Plos One 7(8):e43450.

Ducrocq S, Madur D, Veyrieras J-B, Camus-Kulandaivelu L, Kloiber-Maitz M, Presterl T, Ouzunova M, Manicacci D, Charcosset A. 2008. Key impact of Vgt1 on flowering time adaptation in maize: Evidence from association mapping and ecogeographical information. Genetics 178:2433–2437.

Dudley JW, Lambert RJ. 1992. 90-generattions of selection for oil and protein content in maize. Maydica 37:81–87.

Durand E, Bouchet S, Bertin P, Ressayre A, Jamin P, Charcosset A, Dillmann C, Tenaillon MI. 2012. Flowering Time in maize: linkage and epistasis at a major effect locus. Genetics 190:190(4):1547–62.

Durand E, Tenaillon MI, Raffoux X, Thepot S, Falque M, Jamin P, Bourgais A, Ressayre A, Dillmann C. 2015. Dearth of polymorphism associated with a sustained response to selection for flowering time in maize. BMC Evolutionary Biology 15:103.

Durand E, Tenaillon MI, Ridel C, Coubriche D, Jamin P, Jouanne S, Ressayre A, Charcosset A, Dillmann C. 2010. Standing variation and new mutations both contribute to a fast response to selection for flowering time in maize inbreds. BMC Evolutionary Biology 10:2.

Gage JL, Jarquin D, Romay C, Lorenz A, Buckler ES, Kaeppler S, Alkhalifah N, Bohn M, Campbell DA, Edwards J, et al. 2017. The effect of artificial selection on phenotypic plasticity in maize. Nature Communications 8:1348.

Gervasi DDL, Schiestl FP. 2017. Real-time divergent evolution in plants driven by pollinators. Nature Communications 8:14691.

Goldringer I, Prouin C, Rousset M, Galic N, Bonnin I. 2006. Rapid differentiation of experimental populations of wheat for heading time in response to local climatic conditions. Annals of Botany 98:805–817.

Good BH, McDonald MJ, Barrick JE, Lenski RE, Desai MM. 2017. The dynamics of molecular evolution over 60,000 generations. Nature 551:45–50.

Graves JL, Jr., Hertweck KL, Phillips MA, Han MV, Cabral LG, Barter TT, Greer LF, Burke MK, Mueller LD, Rose MR. 2017. Genomics of Parallel Experimental Evolution in Drosophila. Molecular Biology and Evolution 34:831–842.

Guo L, Wang X, Zhao M, Huang C, Li C, Li D, Yang CJ, York AM, Xue W, Xu G, et al. 2018. Stepwise cis-regulatory changes in ZCN8 contribute to maize flowering-time adaptation. Current biology:28(18):3005–3015.

Hung H-Y, Shannon LM, Tian F, Bradbury PJ, Chen C, Flint-Garcia SA, McMullen MD, Ware D, Buckler ES, Doebley JF, et al. 2012. ZmCCT and the genetic basis of day-length adaptation underlying the postdomestication spread of maize. Proceedings of the National Academy of Sciences of the United States of America 109:E1913–E1921.

Kumar I, Swaminathan K, Hudson K, Hudson ME. 2016. Evolutionary divergence of phytochrome protein function in Zea mays PIF3 signaling. Journal of Experimental Botany 67:4231–4240.

Laehnemann D, Pena-Miller R, Rosenstiel P, Beardmore R, Jansen G, Schulenburg H. 2014. Genomics of Rapid Adaptation to Antibiotics: Convergent Evolution and Scalable Sequence Amplification. Genome Biology and Evolution 6:1287–1301.

Lamkey KR. 1992. 50 years of recurrent selection in the Iowa Stiff Stalk Synthetic maize population. Maydica 37:19–28.

Lehermeier C, Kraemer N, Bauer E, Bauland C, Camisan C, Campo L, Flament P, Melchinger AE, Menz M, Meyer N, et al. 2014. Usefulness of multiparental populations of maize (Zea mays L.) for genome-based prediction. Genetics 198(1):3–16.

Li L, Petsch K, Shimizu R, Liu S, Xu WW, Ying K, Yu J, Scanlon MJ, Schnable PS, Timmermans MCP, et al. 2013. Mendelian and non-mendelian regulation of gene expression in maize. Plos Genetics 14(2): e1007234.

Lopez-Reynoso JD, Hallauer AR. 1998. Twenty-seven cycles of divergent mass selection for ear length in maize. Crop Science 38:1099–1107.

Lorant A, Ross-Ibarra J M.I. T. 2018. Genomics of long- and short-term adaptation in maize and teosinte. PeerJ Preprints 6:e27190v1.

Love MI, Huber W, Anders S. 2014. Moderated estimation of fold change and dispersion for RNA-seq data with DESeq2. Genome Biology 15(12):550.

Maita R, Coors JG. 1996. Twenty cycles of biparental mass selection for prolificacy in the open-pollinated maize population golden glow. Crop Science 36:1527–1532.

Martin M. 2011. Cutadapt removes adapter sequences from high-throughput sequencing reads. EMBnet.journal 17:10–12.

Mascheretti I, Turner K, Brivio RS, Hand A, Colasanti J, Rossi V. 2015. Florigen-encoding genes of day-neutral and photoperiod-sensitive maize are regulated by different chromatin modifications at the floral transition. Plant Physiology 168:1351–U1463.

McGrath PT, Xu Y, Ailion M, Garrison JL, Butcher RA, Bargmann CI. 2011. Parallel evolution of domesticated Caenorhabditis species targets pheromone receptor genes. Nature 477:321–U392.

Meng X, Muszynski MG, Danilevskaya ON. 2011. The FT-like ZCN8 Gene functions as a floral activator and is involved in photoperiod sensitivity in maize. Plant Cell 23:942–960.

Miller TA, Muslin EH, Dorweiler JE. 2008. A maize CONSTANS-like gene, conz1, exhibits distinct diurnal expression patterns in varied photoperiods. Planta 227:1377–1388.

Moose SP, Dudley JW, Rocheford TR. 2004. Maize selection passes the century mark: a unique resource for 21st century genomics. Trends in Plant Science 9:358–364.

Muszynski MG, Dam T, li BL, Shirbroun DM, Hou Z, Bruggemann E, Archibald R, Ananiev EV, Danilevskaya ON. 2006. Delayed flowering1 encodes a basic leucine zipper protein that mediates floral inductive signals at the shoot apex in maize. Plant Physiology 142:1523–1536.

Navarro JAR, Wilcox M, Burgueno J, Romay C, Swarts K, Trachsel S, Preciado E, Terron A, Delgado HV, Vidal V, et al. 2017. A study of allelic diversity underlying flowering-time adaptation in maize landraces (vol 49, pg 476, 2017). Nature Genetics 49:970–970.

Richards CL, Rosas U, Banta J, Bhambhra N, Purugganan MD. 2012. Genome-Wide Patterns of Arabidopsis Gene Expression in Nature. Plos Genetics 8:482–495.

Rodriguez-Verdugo A, Tenaillon O, Gaut BS. 2016. First-Step Mutations during Adaptation Restore the Expression of Hundreds of Genes. Molecular Biology and Evolution 33:25–39.

Roels SAB, Kelly JK. 2011. Rapid evolution caused by pollinator loss in Mimulus guttatus. Evolution 65:2541–2552.

Rohart F, Gautier B, Singh A, Le Cao K-A. 2017. mixOmics: An R package for ’omics feature selection and multiple data integration. Plos Computational Biology 13.

Sakamoto T, Kobayashi M, Itoh H, Tagiri A, Kayano T, Tanaka H, Iwahori S, Matsuoka M. 2001. Expression of a gibberellin 2-oxidase gene around the shoot apex is related to phase transition in rice. Plant Physiology 125:1508–1516.

Salvi S, Sponza G, Morgante M, Tomes D, Niu X, Fengler KA, Meeley R, Ananiev EV, Svitashev S, Bruggemann E, et al. 2007. Conserved noncoding genomic sequences associated with a flowering-time quantitative trait locus m maize. Proceedings of the National Academy of Sciences of the United States of America 104:11376–11381.

Satoh-Nagasawa N, Nagasawa N, Malcomber S, Sakai H, Jackson D. 2006. A trehalose metabolic enzyme controls inflorescence architecture in maize. Nature 441:227–230.

Sekhon RS, Lin HN, Childs KL, Hansey CN, Buell CR, de Leon N, Kaeppler SM. 2011. Genome-wide atlas of transcription during maize development. Plant Journal 66:553–563.

Sheehan MJ, Farmer PR, Brutnell TP. 2004. Structure and expression of maize phytochrome family homeologs. Genetics 167:1395–1405.

Spor A, Kvitek DJ, Nidelet T, Martin J, Legrand J, Dillmann C, Bourgais A, de Vienne D, Sherlock G, Sicard D. 2014. Phenotypic and genotypic convergences are influenced by historical contingency and environment in yeast. Evolution 68:772–790.

Stelpflug SC, Sekhon RS, Vaillancourt B, Hirsch CN, Buell CR, de Leon N, Kaeppler SM. 2016. An expanded maize gene expression atlas based on RNA sequencing and its use to explore root development. Plant Genome 9(1).

Swanson-Wagner R, Briskine R, Schaefer R, Hufford MB, Ross-Ibarra J, Myers CL, Tiffin P, Springer NM. 2012. Reshaping of the maize transcriptome by domestication. Proceedings of the National Academy of Sciences of the United States of America 109:11878–11883.

Takacs EM, Li J, Du C, Ponnala L, Janick-Buckner D, Yu J, Muehlbauer GJ, Schnable PS, Timmermans MCP, Sun Q, et al. 2012. Ontogeny of the maize shoot apical meristem. Plant Cell 24:3219–3234.

Teixeira JEC, Weldekidan T, de Leon N, Flint-Garcia S, Holland JB, Lauter N, Murray SC, Xu W, Hessel DA, Kleintop AE, et al. 2015. Hallauer’s Tuson: a decade of selection for tropical-to-temperate phenological adaptation in maize. Heredity 114:229–240.

Tenaillon O, Rodriguez-Verdugo A, Gaut RL, McDonald P, Bennett AF, Long AD, Gaut BS. 2012. The molecular diversity of adaptive convergence. Science 335:457–461.

Thornsberry JM, Goodman MM, Doebley J, Kresovich S, Nielsen D, Buckler ES. 2001. Dwarf8 polymorphisms associate with variation in flowering time. Nature Genetics 28:286–289.

Trapnell C, Pachter L, Salzberg SL. 2009. TopHat: discovering splice junctions with RNA-Seq. Bioinformatics 25:1105–1111.

Turner CB, Marshall CW, Cooper VS. 2018. Parallel genetic adaptation across environments differing in mode of growth or resource availability. Evolution letters 2:355–367.

Virlouvet L, Jacquemot M-P, Gerentes D, Corti H, Bouton S, Gilard F, Valot B, Trouverie J, Tcherkez G, Falque M, et al. 2011. The ZmASR1 protein influences branched-chain amino acid biosynthesis and maintains kernel yield in maize under water-limited conditions. Plant Physiology 157:917–936.

Vollbrecht E, Springer PS, Goh L, Buckler ES, Martienssen R. 2005. Architecture of floral branch systems in maize and related grasses. Nature 436:1119–1126.

Yang Q, Li Z, Li WQ, Ku LX, Wang C, Ye JR, Li K, Yang N, Li YP, Zhong T, et al. 2013. CACTA-like transposable element in ZmCCT attenuated photoperiod sensitivity and accelerated the postdomestication spread of maize. Proceedings of the National Academy of Sciences of the United States of America 110:16969–16974.

