## Supplemental figures for "Transcriptomic response to divergent selection for flowering times reveals convergence and key players of the underlying gene regulatory network"

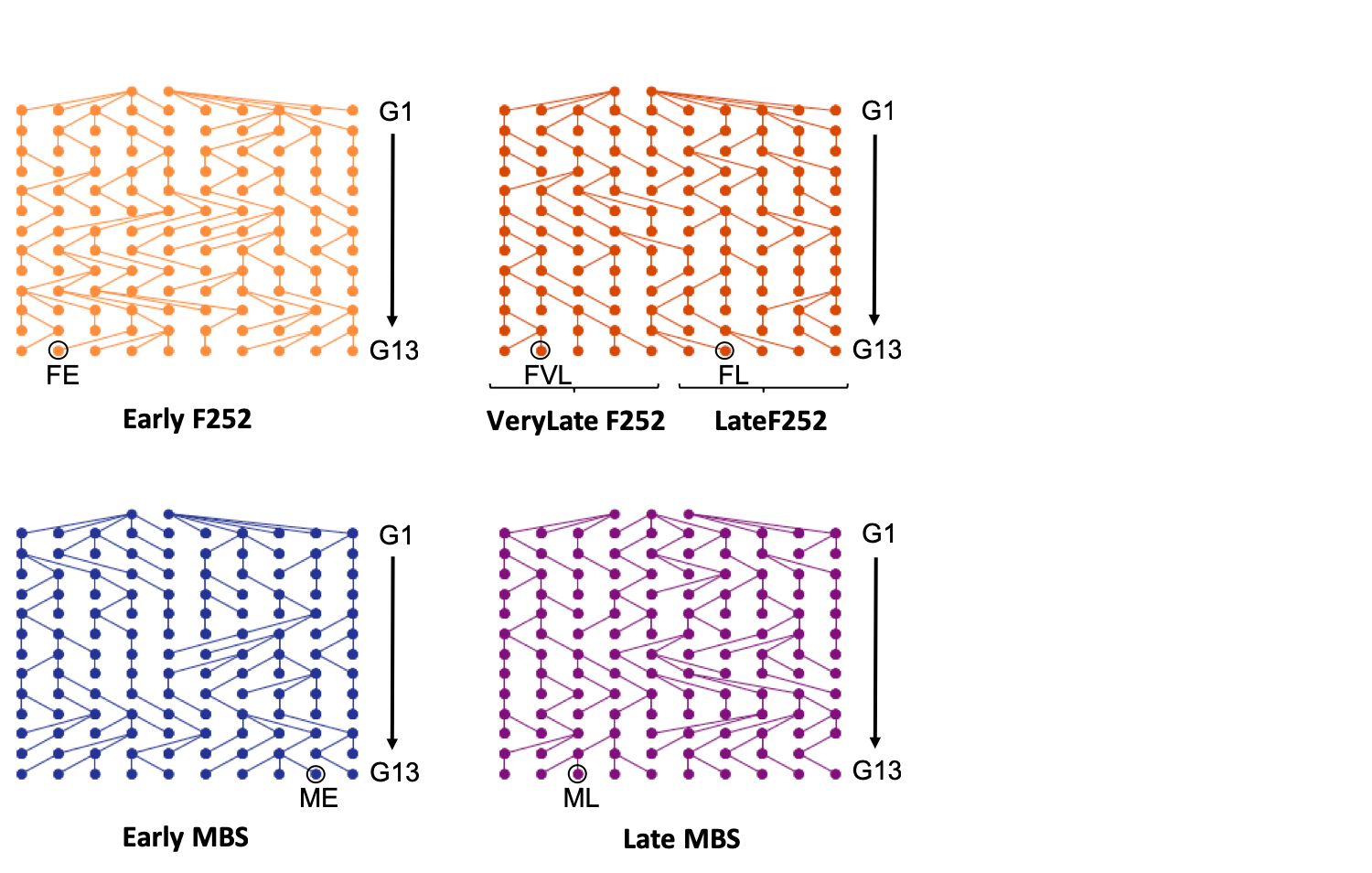

Figure S1. Pedigree of the progenitors in the F252 and MBS Saclay divergent selection experiments (from Durand et al. (2015)). Dots represent selected progenitors from generation G0 to generation G13. We studied 5 progenitors (indicated by black circles) sampled at G13: one Early (FE), one Late(FL) and one VeryLate (FVL) progenitor from F252, as well as one Early (ME) and one Late (ML) progenitor from MBS.

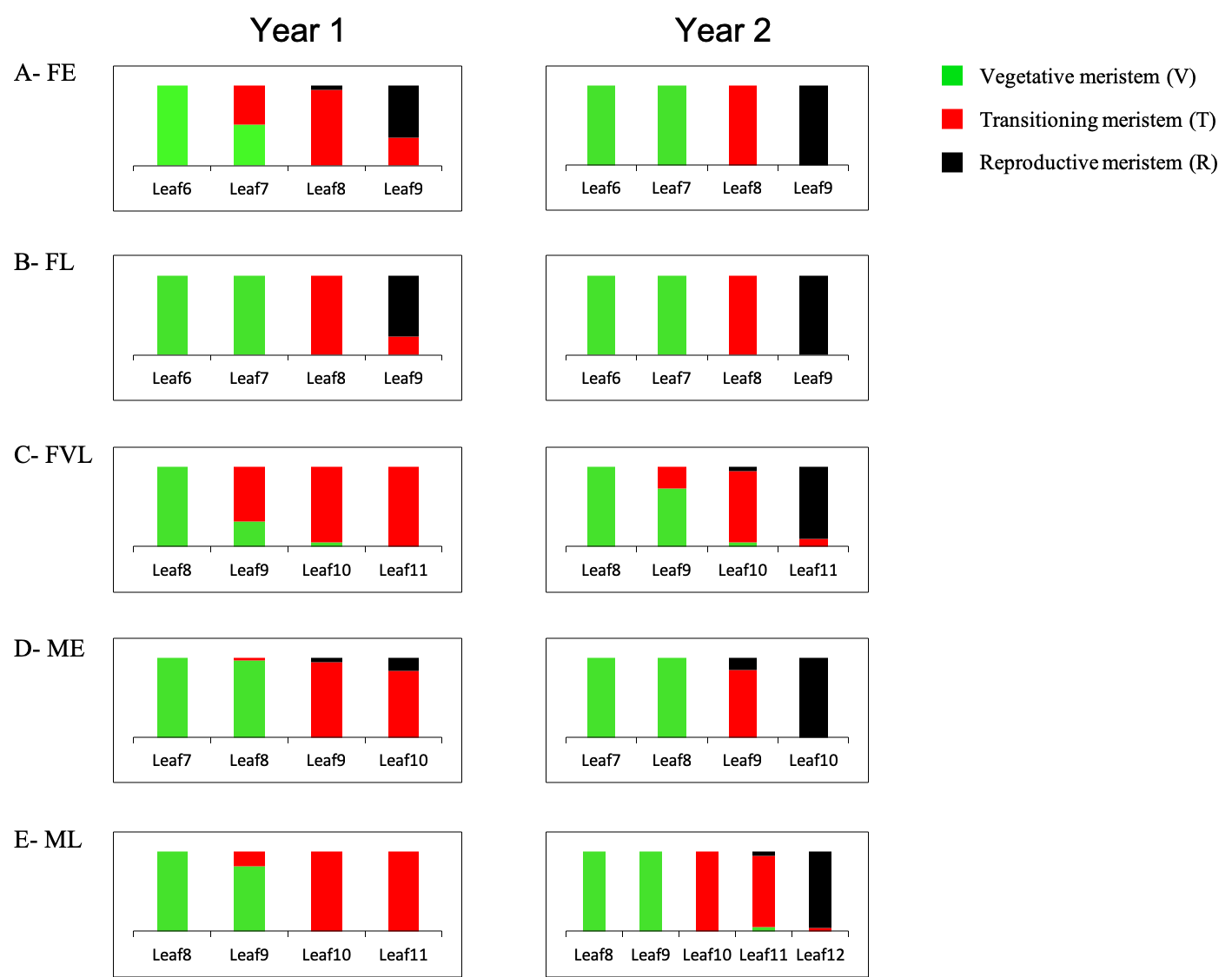

Figure S2. Proportion of Status among all dissected meristems (Table 1) in pooled samples for Year 1 and Year 2 by Progenitor FE (A), FL (B), FVL (C), ME (D), ML (E). Developmental stages are defined as the number (n) of visible leaves.

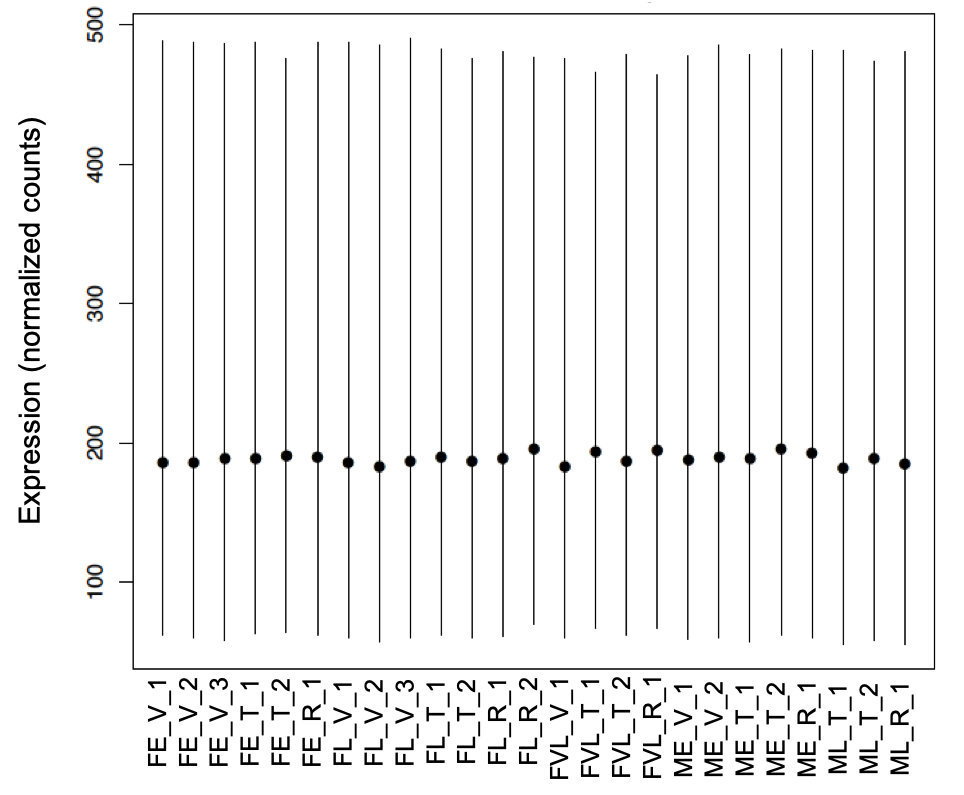

Figure S3. Distribution of normalized counts with median values (dots), 25 and 75 quantiles (vertical lines), for the 25 RNA-seq libraries. Names of the libraries are those described in S1 Table.

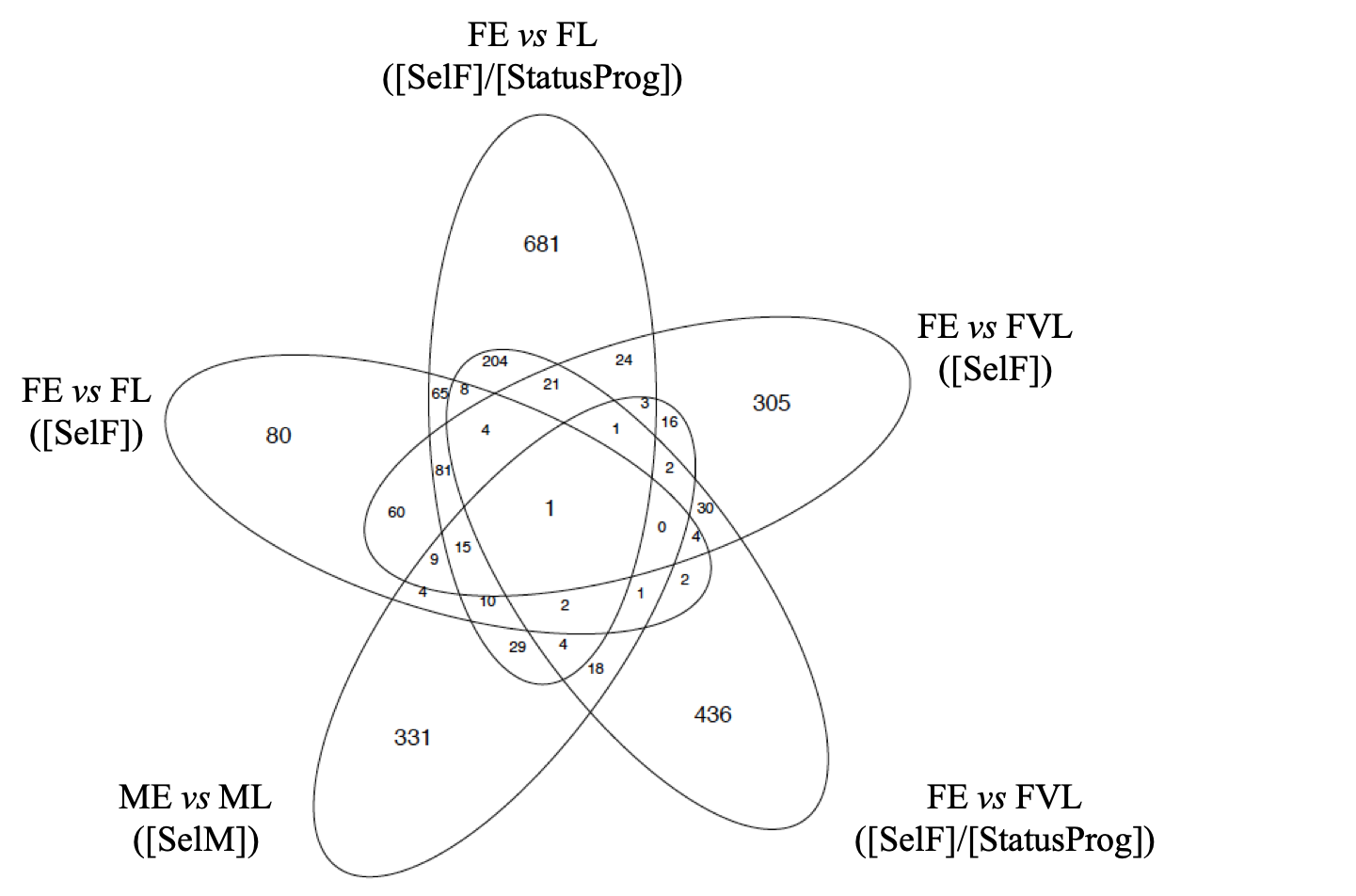

Figure S4. Venn Diagram of DE genes included in the Selection category. The category was further divided into DE genes selected in F252 between FE and FL [SelF], between FE and FVL [SelF], within Status x Progenitor interactions between either FE and FL or FE and FVL [SelF]/[StatusProg], and DE genes selected in MBS [SelM].

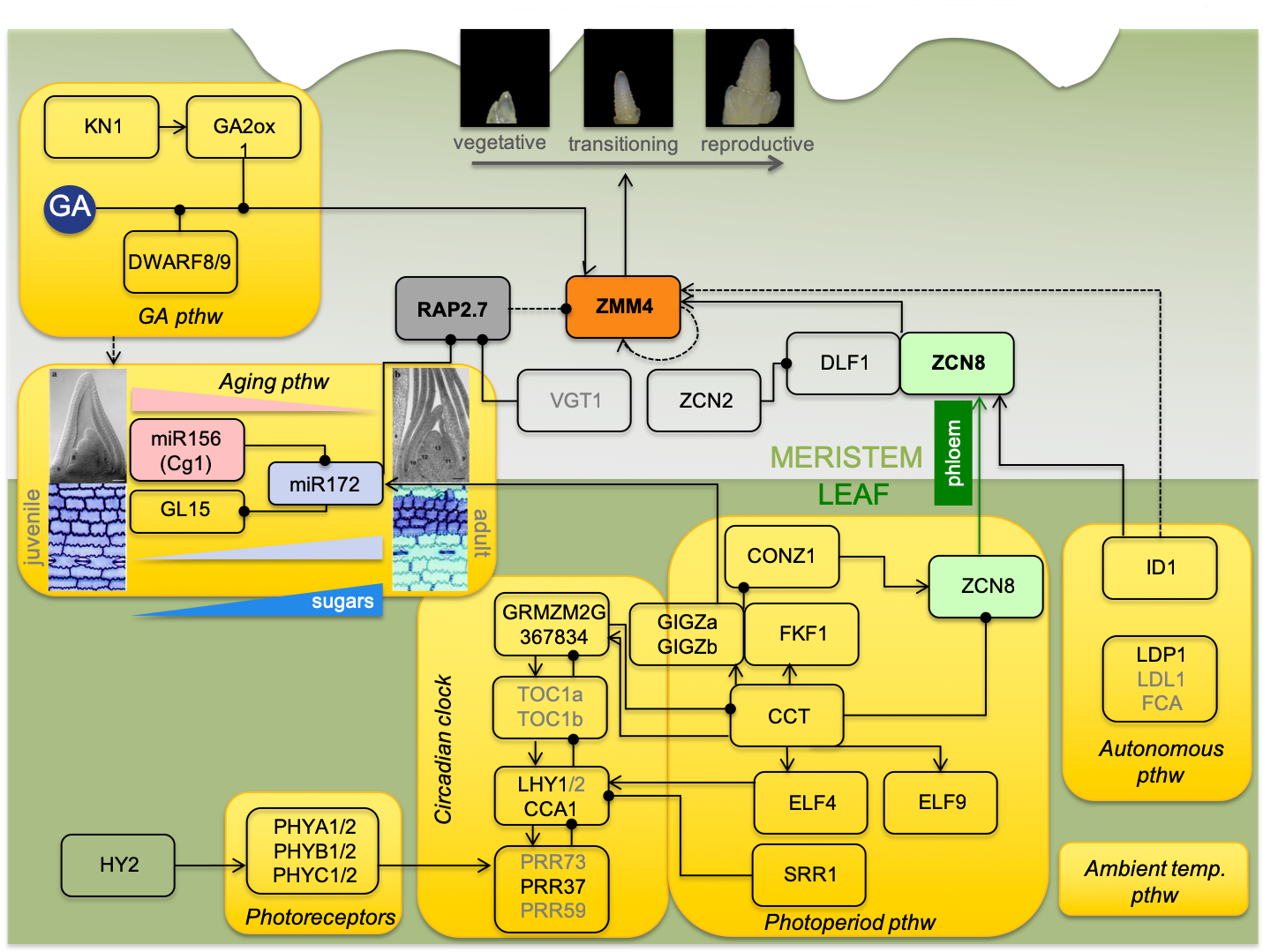

Figure S5. Schematic representation of maize flowering time pathway. Yellow boxes represent different pathways acting in the leaf or the shoot apical meristem or potentially both. Some are well characterized, others not (such as the ambient temperature pathway). When known, relationships between genes are shown with black arrows and dots designating positive and negative regulation respectively. GA designates Gibberellin hormones. Phloem migrates between leaf and the shoot apical meristem transporting flowering signals such as the florigen encoded by *ZCN8*. *ZMM4*, a floral meristem identity integrator, is highlighted in orange as well as a putative negative regulator, *Rap2.7*, highlighted in grey. We investigated the patterns of expression at these three genes in greater details using qRT-PCR. The figure also highlights two miR genes (miR156 in pink and miR172 in purple) with corresponding trends in expression level during plant aging. Putative action of sugar metabolism in floral initiation is shown, and corresponding genes have been added to our flowering candidates. Genes indicated in grey do not display corresponding gene models. Illustration is modified from Dong et al. (2012) with additional information from Lauter et al. (2005), Proveniers et al. (2013), Chuck et al. (2007), Bendix et al. (2013), Yang et al. (2013).

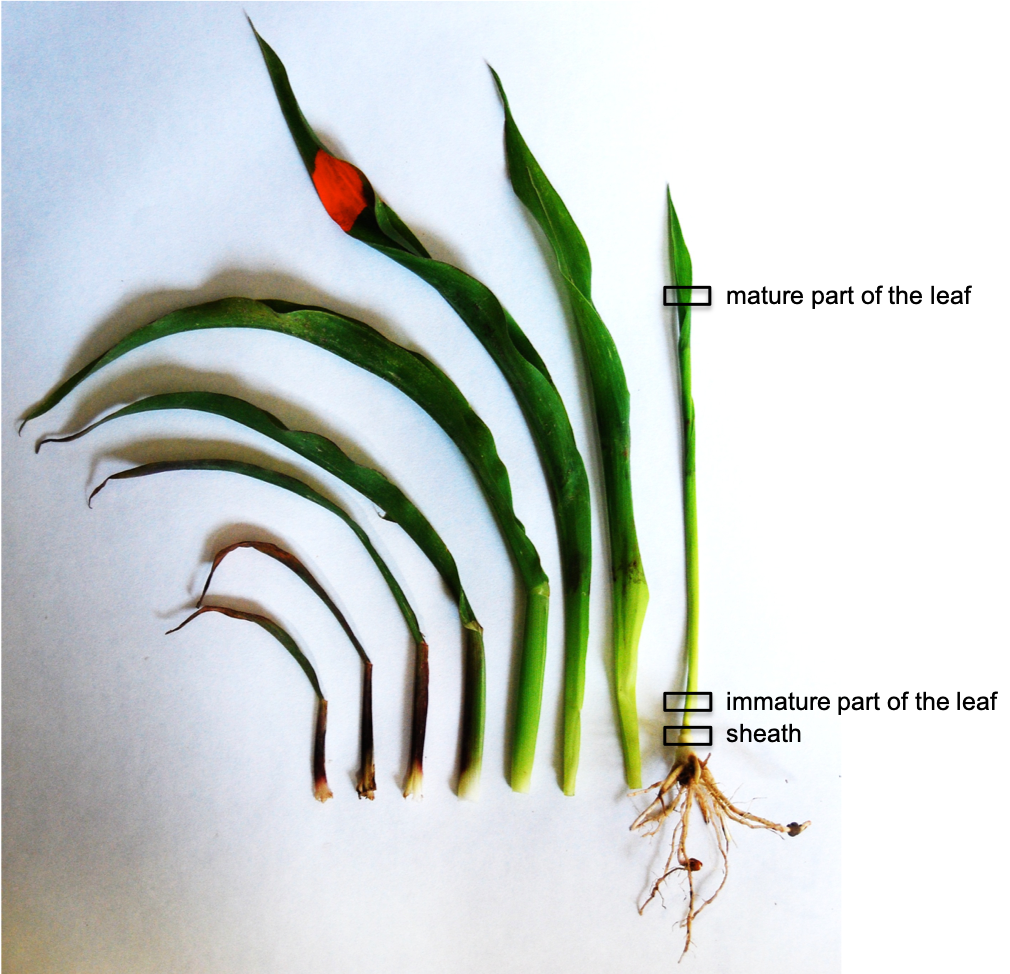

Figure S6. Organs used for the qRT-PCR in addition to the shoot apical meristem. Organs were taken from the last visible leaf, here the Leaf 8.

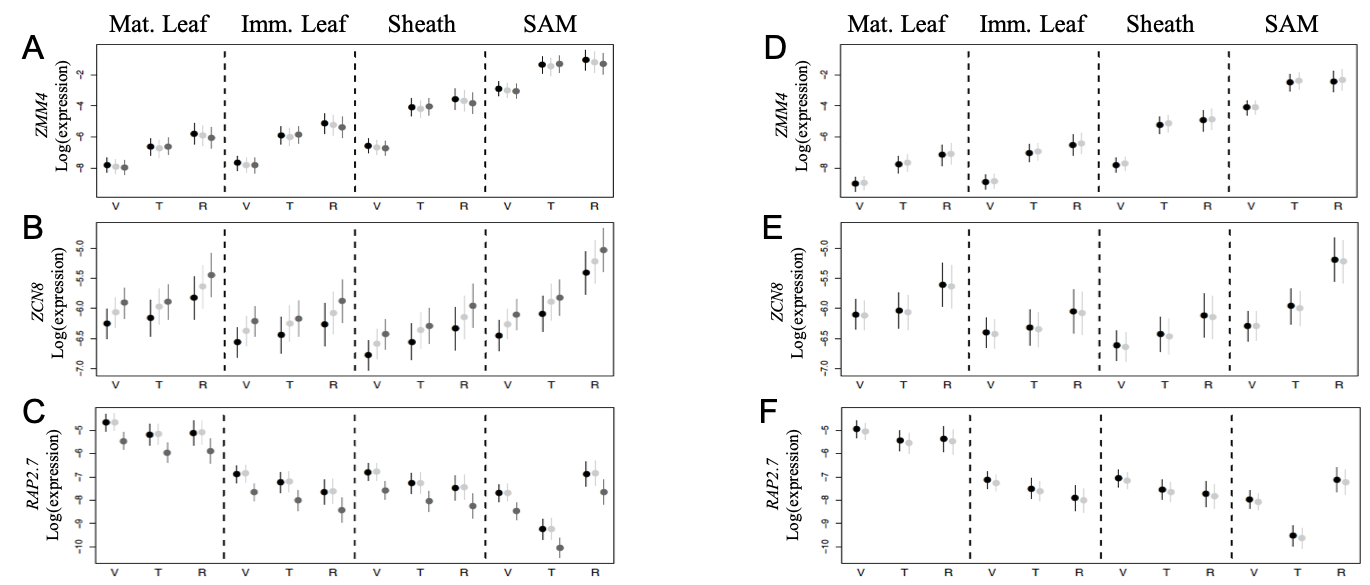

Figure S7. Adjusted means of log(Expression) across Replicates and 95% CI determined by qRT-PCR for 3 genes *ZMM4*, *ZCN8*, *RAP2.7* in F252 (A-C) and MBS (D-F). Expression was determined by shoot apical meristem Status (V=Vegetative, T=Transitioning, R=Reproductive), Organs (mature part, immature part and sheath of the last visible leaf, as well as the shoot apical meristem) and Progenitors (Early, Late, VeryLate from left to right in the F252 panel, and Early, Late from left to right in the MBS panel).

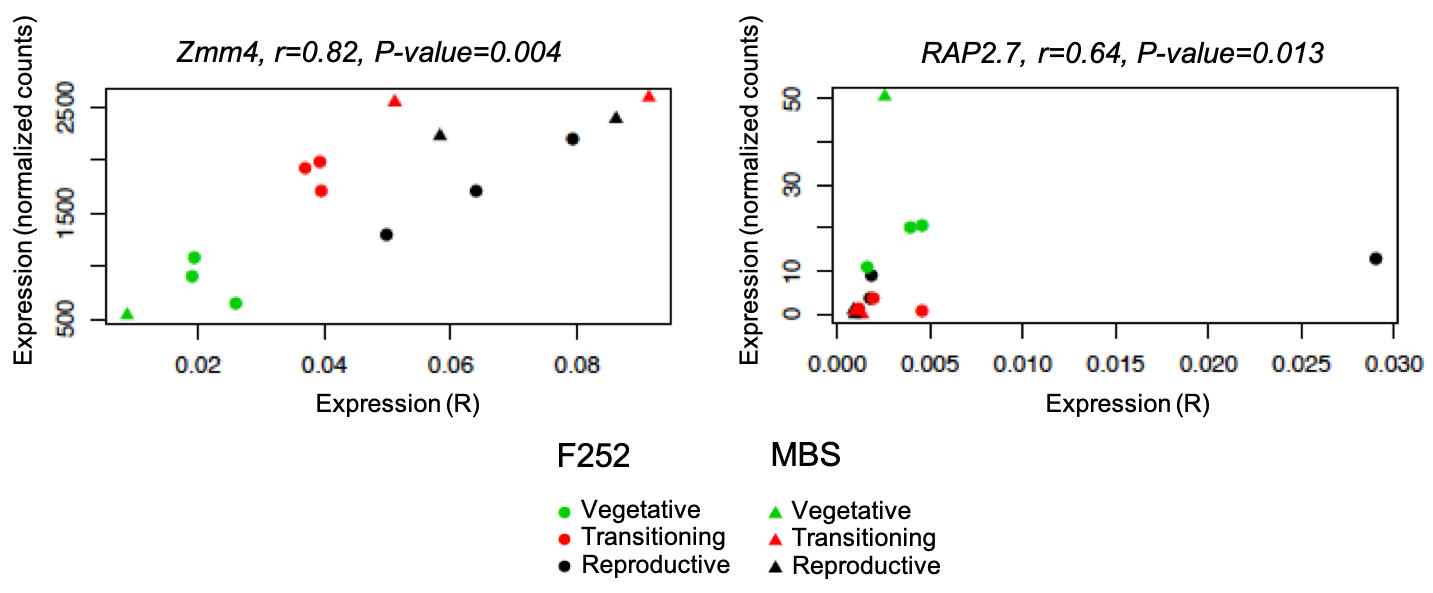

Figure S8. Correlations between levels of expression determined by qRT-PCR and RNA-seq for two candidate genes. Gene expression is expressed as R (ratio of the candidate gene C_T_ over ZmGRP1 C_T_) for qRT-PCR and as normalized read counts for RNA-seq. Spearman correlation coefficients (*r*) were calculated on the common set of 25 samples (RNA-seq libraries) averaged over Progenitors (FE, FL, FVL, ME, ML) and Status (V, T, R). P-values are indicated. We excluded *ZCN8* from analyzes because it displayed no expression in RNA-Seq data.

**Literature cited in Supplementary Figures and Tables**

Alleman M, Sidorenko L, McGinnis K, Seshadri V, Dorweiler JE, White J, Sikkink K, Chandler VL (2006) An RNA-dependent RNA polymerase is required for paramutation in maize. Nature 442: 295-298

Alter P, Bircheneder S, Zhou L-Z, Schlueter U, Gahrtz M, Sonnewald U, Dresselhaus T (2016) Flowering time-regulated genes in maize include the transcription factor ZmMADS1. Plant Physiology 172: 389-404

Bendix C, Mendoza JM, Stanley DN, Meeley R, Harmon FG (2013) The circadian clock-associated gene gigantea1 affects maize developmental transitions. Plant Cell and Environment 36: 1379-1390

Bensen RJ, Johal GS, Crane VC, Tossberg JT, Schnable PS, Meeley RB, Briggs SP (1995) Cloning and characterization of the maize AN1 gene. Plant Cell 7: 75-84

Bolduc N, Hake S (2009) The maize transcription factor KNOTTED1 directly regulates the gibberellin catabolism gene ga2ox1. Plant Cell 21: 1647-1658

Bomblies K, Doebley JF (2006) Pleiotropic effects of the duplicate maize FLORICAULA/LEAFY genes zfl1 and zfl2 on traits under selection during maize domestication. Genetics 172: 519-531

Bommert P, Il Je B, Goldshmidt A, Jackson D (2013) The maize G alpha gene COMPACT PLANT2 functions in CLAVATA signalling to control shoot meristem size. Nature 502: 555-558

Bortiri E, Chuck G, Vollbrecht E, Rocheford T, Martienssen R, Hake S (2006) ramosa2 encodes a lateral organ boundary domain protein that determines the fate of stem cells in branch meristems of maize. Plant Cell 18: 574-585

Buckler ES, Holland JB, Bradbury PJ, Acharya CB, Brown PJ, Browne C, Ersoz E, Flint-Garcia S, Garcia A, Glaubitz JC, Goodman MM, Harjes C, Guill K, Kroon DE, Larsson S, Lepak NK, Li H, Mitchell SE, Pressoir G, Peiffer JA, Rosas MO, Rocheford TR, Cinta Romay M, Romero S, Salvo S, Sanchez Villeda H, da Silva HS, Sun Q, Tian F, Upadyayula N, Ware D, Yates H, Yu J, Zhang Z, Kresovich S, McMullen MD (2009) The genetic architecture of maize flowering time. Science 325: 714-718

Chen Y, Hou M, Liu L, Wu S, Shen Y, Ishiyama K, Kobayashi M, McCarty DR, Tan B-C (2014) The maize DWARF1 encodes a gibberellin 3-oxidase and is dual localized to the nucleus and cytosol. Plant Physiology 166: 2028-2039

Chuck G, Bortiri E (2010) The unique relationship between tsh4 and ra2 in patterning floral phytomers. Plant signaling & behavior 5: 979-981

Chuck G, Cigan AM, Saeteurn K, Hake S (2007) The heterochronic maize mutant Corngrass1 results from overexpression of a tandem microRNA. Nature Genetics 39: 544-549

Chuck G, Meeley R, Hake S (2008) Floral meristem initiation and meristem cell fate are regulated by the maize AP2 genes ids1 and sid1. Development 135: 3013-3019

Chuck G, Meeley R, Irish E, Sakai H, Hake S (2007) The maize tasselseed4 microRNA controls sex determination and meristem cell fate by targeting Tasselseed6/indeterminate spikelet1. Nature Genetics 39: 1517-1521

Chuck G, Whipple C, Jackson D, Hake S (2010) The maize SBP-box transcription factor encoded by tasselsheath4 regulates bract development and the establishment of meristem boundaries. Development 137: 1585-1585

Chuck GS, Brown PJ, Meeley R, Hake S (2014) Maize SBP-box transcription factors unbranched2 and unbranched3 affect yield traits by regulating the rate of lateral primordia initiation. Proceedings of the National Academy of Sciences of the United States of America 111: 18775-18780

Chuck GS, Tobias C, Sun L, Kraemer F, Li C, Dibble D, Arora R, Bragg JN, Vogel JP, Singh S, Simmons BA, Pauly M, Hake S (2011) Overexpression of the maize Corngrass1 microRNA prevents flowering, improves digestibility, and increases starch content of switchgrass. Proceedings of the National Academy of Sciences of the United States of America 108: 17550-17555

Colasanti J, Yuan Z, Sundaresan V (1998) The indeterminate gene encodes a zinc finger protein and regulates a leaf-generated signal required for the transition to flowering in maize. Cell 93: 593-603

Coneva V, Zhu T, Colasanti J (2007) Expression differences between normal and indeterminate1 maize suggest downstream targets of ID1, a floral transition regulator in maize. Journal of Experimental Botany 58: 3679-3693

Danilevskaya ON, Meng X, Ananiev EV (2010) Concerted modification of flowering time and inflorescence architecture by ectopic expression of TFL1-Like genes in maize. Plant Physiology 153: 238-251

Danilevskaya ON, Meng X, Selinger DA, Deschamps S, Hermon P, Vansant G, Gupta R, Ananiev EV, Muszynski MG (2008) Involvement of the MADS-Box gene ZMM4 in floral induction and inflorescence development in maize. Plant Physiology 147: 2054-2069

Dong Z, Danilevskaya O, Abadie T, Messina C, Coles N, Cooper M (2012) A gene regulatory network model for floral transition of the shoot apex in maize and its dynamic modeling. Plos One 7: e43450

Durand E, Bouchet S, Bertin P, Ressayre A, Jamin P, Charcosset A, Dillmann C, Tenaillon MI (2012) Flowering time in maize: Linkage and epistasis at a major effect locus. Genetics 190: 1547-1562

Durand E, Tenaillon MI, Raffoux X, Thepot S, Falque M, Jamin P, Bourgais A, Ressayre A, Dillmann C (2015) Dearth of polymorphism associated with a sustained response to selection for flowering time in maize. BMC Evolutionary Biology 15: 103

Evans MMS, Poethig RS (1995) Gibberellins promote vegetative phase-change and reproductive maturity in maize. Plant Physiology 108: 475-487

Evans MMS, Poethig RS (1997) The viviparous8 mutation delays vegetative phase change and accelerates the rate of seedling growth in maize. Plant Journal 12: 769-779

Gallavotti A, Zhao Q, Kyozuka J, Meeley RB, Ritter M, Doebley JF, Pe ME, Schmidt RJ (2004) The role of barren stalk1 in the architecture of maize. Nature 432: 630-635

Hayes KR, Beatty M, Meng X, Simmons CR, Habben JE, Danilevskaya ON (2010) Maize global transcriptomics reveals pervasive leaf diurnal rhythms but rhythms in developing ears are largely limited to the core oscillator. Plos One 5: e12887

He C, Fu JJ, Zhang J, Li YX, Zheng J, Zhang HW, Yang XH, Wang JH, Wang GY (2017) A gene-oriented haplotype comparison reveals recently selected genomic regions in temperate and tropical maize germplasm. Plos One 12: e0169806

Heuer S, Hansen S, Bantin J, Brettschneider R, Kranz E, Lorz H, Dresselhaus T (2001) The maize MADS box gene ZmMADS3 affects node number and spikelet development and is co-expressed with ZmMADS1 during flower development, in egg cells, and early embryogenesis. Plant Physiology 127: 33-45

Hung H-Y, Shannon LM, Tian F, Bradbury PJ, Chen C, Flint-Garcia SA, McMullen MD, Ware D, Buckler ES, Doebley JF, Holland JB (2012) ZmCCT and the genetic basis of day-length adaptation underlying the postdomestication spread of maize. Proceedings of the National Academy of Sciences of the United States of America 109: E1913-E1921

Kumar I, Swaminathan K, Hudson K, Hudson ME (2016) Evolutionary divergence of phytochrome protein function in Zea mays PIF3 signaling. Journal of Experimental Botany 67: 4231-4240

Lauter N, Kampani A, Carlson S, Goebel M, Moose SP (2005) microRNA172 down-regulates glossy15 to promote vegetative phase change in maize. Proceedings of the National Academy of Sciences of the United States of America 102: 9412-9417

Lawit SJ, Wych HM, Xu D, Kundu S, Tomes DT (2010) Maize DELLA proteins dwarf plant8 and dwarf plant9 as modulators of plant development. Plant and Cell Physiology 51: 1854-1868

Lee B-h, Johnston R, Yang Y, Gallavotti A, Kojima M, Travencolo BAN, Costa LdF, Sakakibara H, Jackson D (2009) Studies of aberrant phyllotaxy1 mutants of maize indicate complex interactions between auxin and cytokinin signaling in the shoot apical meristem. Plant Physiology 150: 205-216

Li X, Zhu C, Yeh C-T, Wu W, Takacs EM, Petsch KA, Tian F, Bai G, Buckler ES, Muehlbauer GJ, Timmermans MCP, Scanlon MJ, Schnable PS, Yu J (2012) Genic and nongenic contributions to natural variation of quantitative traits in maize. Genome Research 22: 2436-2444

Mascheretti I, Turner K, Brivio RS, Hand A, Colasanti J, Rossi V (2015) Florigen-encoding genes of day-neutral and photoperiod-sensitive maize are regulated by different chromatin modifications at the floral transition. Plant Physiology 168: 1351-U1463

McSteen P, Malcomber S, Skirpan A, Lunde C, Wu X, Kellogg E, Hake S (2007) barren inflorescence2 encodes a co-ortholog of the PINOID serine/threonine kinase and is required for organogenesis during inflorescence and vegetative development in maize. Plant Physiology 144: 1000-1011

Meng X, Muszynski MG, Danilevskaya ON (2011) The FT-like ZCN8 gene functions as a floral activator and is involved in photoperiod sensitivity in maize. Plant Cell 23: 942-960

Miller TA, Muslin EH, Dorweiler JE (2008) A maize CONSTANS-like gene, conz1, exhibits distinct diurnal expression patterns in varied photoperiods. Planta 227: 1377-1388

Murphy RL, Klein RR, Morishige DT, Brady JA, Rooney WL, Miller FR, Dugas DV, Klein PE, Mullet JE (2011) Coincident light and clock regulation of pseudoresponse regulator protein 37 (PRR37) controls photoperiodic flowering in sorghum. Proceedings of the National Academy of Sciences of the United States of America 108: 16469-16474

Muszynski MG, Dam T, li BL, Shirbroun DM, Hou Z, Bruggemann E, Archibald R, Ananiev EV, Danilevskaya ON (2006) Delayed flowering1 encodes a basic leucine zipper protein that mediates floral inductive signals at the shoot apex in maize. Plant Physiology 142: 1523-1536

Pautler M, Eveland AL, LaRue T, Yang F, Weeks R, Lunde C, Il Je B, Meeley R, Komatsu M, Vollbrecht E, Sakai H, Jackson D (2015) FASCIATED EAR4 Encodes a bZIP transcription factor that regulates shoot meristem size in maize. Plant Cell 27: 104-120

Proveniers M (2013) Sugars speed up the circle of life. Elife 2

Salvi S, Sponza G, Morgante M, Tomes D, Niu X, Fengler KA, Meeley R, Ananiev EV, Svitashev S, Bruggemann E, Li B, Hainey CF, Radovic S, Zaina G, Rafalski JA, Tingey SV, Miao G-H, Phillips RL, Tuberosa R (2007) Conserved noncoding genomic sequences associated with a flowering-time quantitative trait locus m maize. Proceedings of the National Academy of Sciences of the United States of America 104: 11376-11381

Satoh-Nagasawa N, Nagasawa N, Malcomber S, Sakai H, Jackson D (2006) A trehalose metabolic enzyme controls inflorescence architecture in maize. Nature 441: 227-230

Sawers RJH, Linley PJ, Gutierrez-Marcos JF, Delli-Bovi T, Farmer PR, Kohchi T, Terry MJ, Brutnell TP (2004) The Elm1 (ZmHy2) gene of maize encodes a phytochromobilin synthase. Plant Physiology 136: 2771-2781

Sheehan MJ, Farmer PR, Brutnell TP (2004) Structure and expression of maize phytochrome family homeologs. Genetics 167: 1395-1405

Taguchi-Shiobara F, Yuan Z, Hake S, Jackson D (2001) The fasciated ear2 gene encodes a leucine-rich repeat receptor-like protein that regulates shoot meristem proliferation in maize. Genes & Development 15: 2755-2766

Takacs EM, Li J, Du C, Ponnala L, Janick-Buckner D, Yu J, Muehlbauer GJ, Schnable PS, Timmermans MCP, Sun Q, Nettleton D, Scanlon MJ (2012) Ontogeny of the maize shoot apical meristem. Plant Cell 24: 3219-3234

Teotia S, Tang GL (2015) To bloom or not to bloom: role of microRNAs in plant flowering. Molecular Plant 8: 359-377

Thornsberry JM, Goodman MM, Doebley J, Kresovich S, Nielsen D, Buckler ES (2001) Dwarf8 polymorphisms associate with variation in flowering time. Nature Genetics 28: 286-289

van Nocker S, Muszynski M, Briggs K, Amasino RM (2000) Characterization of a gene from *Zea mays* related to the *Arabidopsis* flowering-time gene LUMINIDEPENDENS. Plant Molecular Biology 44: 107-122

Vega SH, Sauer M, Orkwiszewski JAJ, Poethig RS (2002) The early phase change gene in Maize. Plant Cell 14: 133-147

Vollbrecht E, Springer PS, Goh L, Buckler ES, Martienssen R (2005) Architecture of floral branch systems in maize and related grasses. Nature 436: 1119-1126

Wang X, Wu L, Zhang S, Wu L, Ku L, Wei X, Xie L, Chen Y (2011) Robust expression and association of ZmCCA1 with circadian rhythms in maize. Plant Cell Reports 30: 1261-1272

Whipple CJ, Kebrom TH, Weber AL, Yang F, Hall D, Meeley R, Schmidt R, Doebley J, Brutnell TP, Jackson DP (2011) grassy tillers1 promotes apical dominance in maize and responds to shade signals in the grasses. Proceedings of the National Academy of Sciences of the United States of America 108: E506-E512

Woodward JB, Abeydeera ND, Paul D, Phillips K, Rapala-Kozik M, Freeling M, Begley TP, Ealick SE, McSteen P, Scanlon MJ (2010) A maize thiamine auxotroph is defective in shoot meristem maintenance. Plant Cell 22: 3305-3317

Yang Q, Li Z, Li WQ, Ku LX, Wang C, Ye JR, Li K, Yang N, Li YP, Zhong T, Li JS, Chen YH, Yan JB, Yang XH, Xu ML (2013) CACTA-like transposable element in ZmCCT attenuated photoperiod sensitivity and accelerated the postdomestication spread of maize. Proceedings of the National Academy of Sciences of the United States of America 110: 16969-16974
